# LuCEMe: An accessible luciferase-based method to functionally assess cellular metabolism across model systems

**DOI:** 10.64898/2026.11.07.765041

**Authors:** Lotfi Laaouimir, Célia Sergi, Nick van Gastel

## Abstract

Metabolic profiling approaches such as extracellular flux analysis, metabolomics, stable isotope tracing and cytometry-based assays have transformed our understanding of cellular metabolism in physiological and pathological contexts. However, most techniques require specialized equipment or reagents, and do not allow differential metabolic analysis in intact multicellular systems. Here, we present LuCEMe, a luciferase-based metabolic phenotyping approach that combines acute metabolic perturbations with quantitative ATP measurements to resolve glycolytic and mitochondrial dependencies and capacities under native culture conditions, including in complex coculture and ex vivo systems without prior cell separation. The assay can also be adapted to increase metabolic resolution and distinguish between glycolytic activity, glucose oxidation and fatty acid/amino acid-driven mitochondrial ATP production. We use LuCEMe to reveal profound metabolic differences between leukemia cells cocultured with macrophages, stromal cells or adipocytes, and demonstrate distinct metabolic rewiring during macrophage polarization, osteogenic and adipogenic differentiation, and T cell activation. Ex vivo LuCEMe analysis identified a particular metabolic adaptation associated with chemotherapy response in acute myeloid leukemia that was not evident from conventional analyses. LuCEMe thus provides an accessible, robust and versatile framework for direct functional metabolic phenotyping across in vitro, ex vivo, and multicellular systems that complements existing metabolic profiling technologies.

## Introduction

Cellular energy metabolism is a highly adaptive process that enables cells to meet energetic and biosynthetic demands in response to fluctuations in nutrient availability, extracellular signals, and cell-intrinsic regulatory programs. Through the coordinated control of glycolytic flux, mitochondrial oxidative metabolism and associated metabolic pathways, cells dynamically modulate ATP production to support essential cellular functions, including survival, proliferation, and differentiation^1^. In cancer, these regulatory mechanisms are frequently altered, resulting in profound metabolic reprogramming that supports malignant growth and persistence. Alterations in glucose metabolism and mitochondrial oxidative phosphorylation (OXPHOS) are among the most prominent features of this reprogramming^2^. Consequently, defining the metabolic pathways and substrates that sustain cellular ATP generation is fundamental to understanding both physiological and pathological states.

Metabolomics techniques, including mass spectrometry- or nuclear magnetic resonance spectroscopy-based methods, provide a global overview of the cellular metabolome but require large cell numbers and advanced equipment for analysis, and do not give functional insight. The latter can be addressed through stable isotope tracing and metabolic flux analysis, but this further increases the need for specialized reagents, analysis pipelines and scientific expertise, limiting broad implementation. To address this need, several experimental approaches have been developed to functionally probe cellular bioenergetics in a more accessible manner, among which extracellular flux analysis and protein synthesis-based metabolic dependency assays are widely used and have substantially advanced the field. Extracellular flux-based technologies, such as Seahorse XF analysis, enable real-time measurements of oxygen consumption rate (OCR) and extracellular acidification rate (ECAR), providing important insights into mitochondrial respiratory function, glycolytic engagement, and bioenergetic flexibility^3,4^. However, Seahorse assays typically involve extended preparatory steps, including cell replating, equilibration in assay-specific media, and sequential pharmacological perturbations. Given the rapid adaptability of metabolic networks, these manipulations may influence cellular metabolic states prior to or during measurement, complicating assessment of intrinsic energetic dependencies.

Complementary approaches such as SCENITH allow to estimate pathway reliance by quantifying protein synthesis under acute inhibition of glycolysis or the mitochondrial electron transport chain using respectively 2-deoxy-D-glucose (2DG) and oligomycin (OG) ^5^. This strategy enables pathway-level comparisons across diverse cell types without requiring specialized metabolic instrumentation and has proven particularly useful in immunological and heterogeneous cellular contexts. An important advantage of this approach is its compatibility with limited cell numbers and analysis of rare cellular subsets, enabling metabolic interrogation of primary samples and phenotypically defined populations that may not be amenable to bulk metabolic assays. The ability to combine energetic profiling with high-dimensional immunophenotyping has established SCENITH as a valuable platform for dissecting metabolic heterogeneity across complex tissues and disease settings^5^. Nevertheless, SCENITH infers metabolic activity indirectly by measuring protein translation, which is influenced not only by ATP availability but also by stress signaling. For instance, 2DG not only interferes with glycolysis, but it can also block N-linked glycosylation due to its structural similarity to mannose, leading to ER stress and unfolded protein response that impact translation^6^. Furthermore, OG-mediated inhibition of complex V can lead to DELE1 cytosolic accumulation which results in the phosphorylation of elF2a, thus blocking protein synthesis^7^. Interpretation using this approach can therefore be influenced by assumptions linking protein synthesis rates to ATP availability.

An additional consideration for existing functional metabolic assays is their application to multicellular or in vitro coculture systems. Many metabolic assays are optimized for relatively homogeneous cell populations, whereas cell-cell interactions, nutrient competition, and paracrine signaling are critical determinants of metabolic behavior. Dissecting cell type-specific metabolic requirements in cocultures remains challenging, as extracellular flux analysis or metabolomics techniques require physical separation of cell types prior to analysis. Given that SCENITH uses flow cytometry for readout, performing it on coculture settings is possible. However, detaching cells from culture plates after coculture requires the usage of enzymes such as trypsin and collagenase which can influence cell metabolism prior to the metabolic assay^8–10^.

Taken together, existing metabolic assays provide powerful and complementary insights into cellular bioenergetics, yet each of the existing techniques comes with certain limitations. In this study, we present a rapid and scalable functional metabolic assay that directly assesses metabolic pathways sustaining cellular ATP synthesis over short timescales, without the disruption of (co)culture conditions for analysis, thereby reducing manipulation-induced confounding effects. We took advantage of the luciferase enzyme that requires ATP to catalyze the oxidation of its substrate luciferin, and thereby emitting light, as a direct readout of ATP levels after exposure to different metabolic challenges. This allows to directly measure metabolic dependencies and capacities through bioluminescence signal measurement. By using either a commercially available kit or by endogenously expressing luciferase in various cells of interest or in different cellular compartments, this technique offers a flexible and robust metabolic readout that effectively democratizes access to metabolic phenotyping. We termed this method LuCEMe for Luciferase-based Cellular Energy Metabolism analysis. We validated the LuCEMe assay using primary mouse cells and various cell lines in mono- or coculture setups, thus introducing a new, simple and accessible method for the analysis of cellular metabolism.

## Design

To enable rapid metabolic phenotyping of cells while preserving native culture conditions, we designed an assay that directly quantifies the metabolic sources of ATP through acute pathway perturbation. The overall strategy is to measure short-timescale changes in intracellular ATP levels following selective inhibition of defined bioenergetic pathways, thereby functionally assigning ATP generation dependencies and capacities without reliance on surrogate readouts. Cells that are either expressing luciferase or not are exposed to inhibitors targeting glycolysis (2DG), OXPHOS (OG), or both, after which intracellular ATP is quantified immediately by either the addition of luciferin to the culture plates and then measuring luminescence in real time, or by using a commercial luciferase-based ATP quantification assay as an endpoint readout. Because the assay quantifies ATP itself, it captures the integrated output of all ATP-producing reactions while preserving pathway specificity through controlled perturbation (Figure 1A).

**Figure 1:**
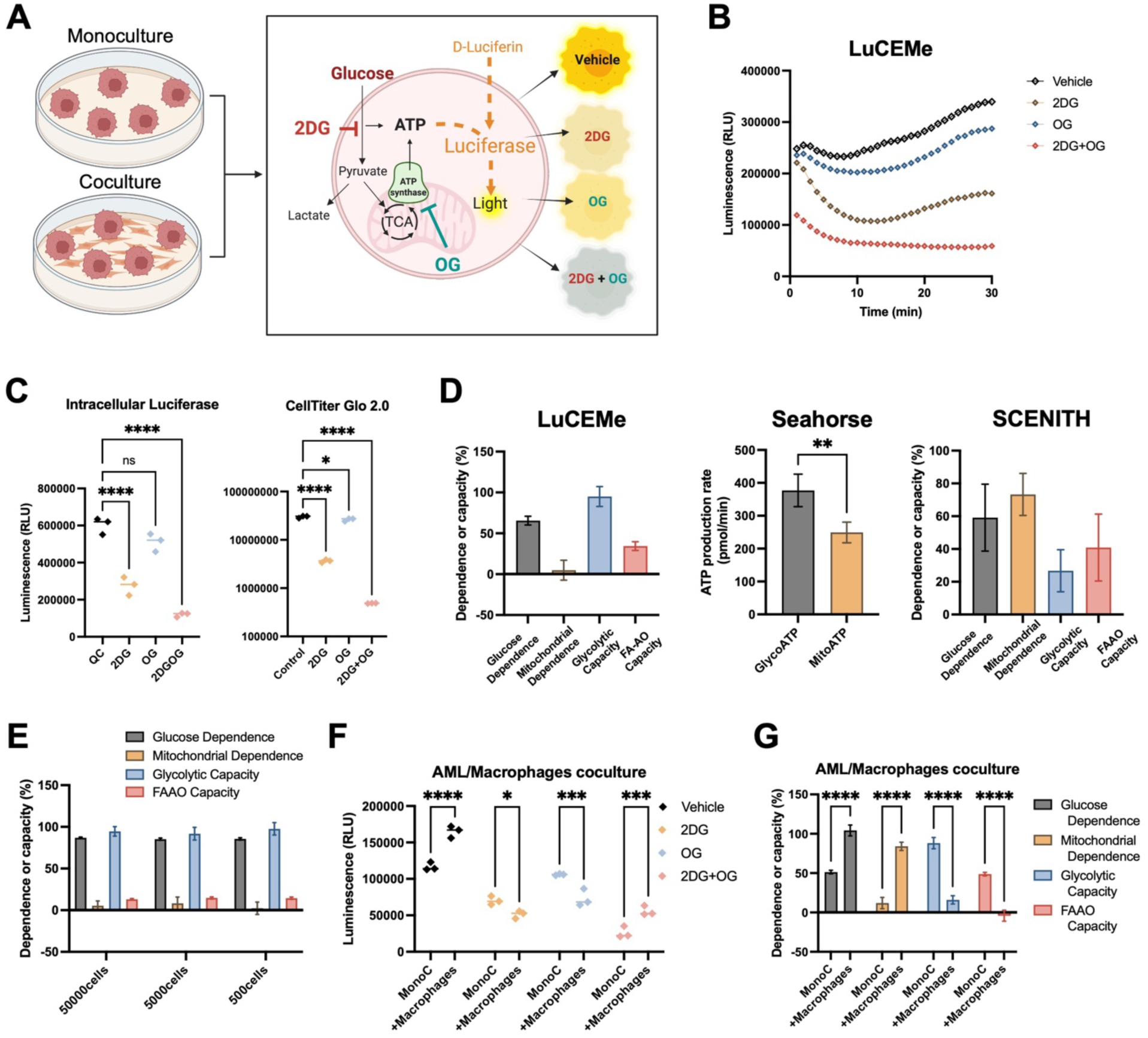
LuCEMe design and proof of concept. **(A)** Schematic overview of the LuCEMe experimental design. 2DG: 2-deoxy-D-glucose; OG: oligomycin. **(B)** Visualization of fluctuations of ATP levels measured in real time after blocking glycolysis and mitochondrial metabolism in MLL-AF9 AML cells expressing luciferase. **(C)** Comparison of ATP levels measured using intracellular luciferase or using CellTiter Glo on AML cells after exposure to 2DG and/or OG for 30 minutes. **(D)** The metabolic profile of AML cells after LuCEMe, Seahorse and SCENITH analyses. **(E)** The metabolic profile of AML cells seeded at different cell numbers determined by LuCEMe. **(F)** ATP levels of AML cells in coculture with macrophages for 24 hours after exposure to 2DG and/or OG for 30 minutes. **(G)** The metabolic profile of AML cells in monoculture or in coculture with macrophages for 24 hours determined by LuCEMe. Data are represented as mean ± standard deviation (SD). *p < 0.05, **p < 0.01, ***p < 0.001.

To test our method, we used a mouse acute myeloid leukemia (AML) cell line carrying the MLL-AF9 translocation that constitutively and ubiquitously expresses intracellular luciferase^11^. AML cells are known to exhibit context-dependent changes in glucose utilization alongside a sustained reliance on OXPHOS^2,11,12^. We detected an immediate drop of ATP after exposure to 2DG, OG and an even more pronounced drop after exposure to the combination. Interestingly, after 10 minutes of exposure to the metabolic inhibitors, ATP levels slightly increased in both the 2DG and OG conditions but not in the combination, suggesting metabolic compensation to retain ATP levels by upregulating respectively OXPHOS or glycolysis (Figure 1B). The same changes in ATP levels after exposure to 2DG, OG or the combination was detected in these AML cells using CellTiter Glo as an endpoint readout, confirming that LuCEMe can be performed using a commercially available kit (Figure 1C). The magnitude and kinetics of ATP decline under each condition define the relative contribution of specific pathways to basal ATP maintenance, which can be used to calculate metabolic dependencies and capacities (Figure 1D). With this setup we were able to see that the AML cells rely mostly on glucose to generate ATP, with a substantial glycolytic capacity compared to their fatty acid/amino acid oxidation (FAAO) capacity. We then compared the results of our LuCEMe assay to existing techniques. Using Seahorse XF analysis, a similar phenotype was detected, where the highest proportion of ATP is generated from glycolysis in these AML cells (Figure 1D). Interestingly, this metabolic profile was not observed when SCENITH was used as a readout, as protein synthesis was similarly decreased by OG and 2DG, which indicates a more pronounced mitochondrial dependence (Figure 1D, Supplemental Figure 1A). The discrepancy between the results obtained using LuCEMe and Seahorse XF on one hand and SCENITH on the other could be due to effects of mitochondrial complex V inhibition on protein synthesis independent of ATP levels, or unspecific binding of the 12D10 anti-puromycin antibody (as clone R4743L-E8 used in the original SCENITH study is not commercially available) ^5–7^. As SCENITH allows metabolic profiling of rare and low-input cell populations, we next assessed whether LuCEMe could similarly preserve metabolic measurements with low cell numbers. For that, AML cells were seeded at progressively lower densities and analyzed under identical experimental conditions. Remarkably, reducing the number of input cells from 50 000 to 5 000 or 500 did not alter the metabolic phenotype, as glucose dependence, mitochondrial dependence, glycolytic capacity, and FAAO capacity remained stable across all cell densities tested (Figure 1E, Supplemental Figure 1B). These results demonstrate that the assay robustly captures cellular metabolic dependencies even from low-input samples, thereby extending its applicability to experimental settings in which cell availability is limiting.

A key design objective was compatibility with multicellular and coculture models. In contrast to approaches that require post-culture physical separation, LuCEMe can be performed in intact cocultures if luciferase is expressed by only one cell type and can thus be applied to adherent-suspension cocultures, stromal-tumor systems or immune-target cell interactions while preserving cell-cell contacts and local nutrient gradients. To confirm this, we cocultured the same MLL-AF9 AML mouse cell line expressing luciferase with a bone marrow macrophage cell line for 24 hours and then exposed the cocultures to either 2DG, OG or the combination. Luciferin was added immediately, and luminescence was measured in real time. We observed that AML cells in coculture with macrophages had higher ATP levels at baseline but showed a more pronounced decrease of ATP after exposure to either 2DG or OG while retaining more ATP after exposure to the combination of 2DG and OG (Figure 1F). With this set-up, we calculated a significant increase of both AML cells’ glucose and mitochondrial dependence and a decrease of their glycolytic and their FAAO capacities after coculture with macrophages for 24 hours (Figure 1G). Based on these results we conclude that upon coculture with macrophages, MLL-AF9 AML cells shift their metabolism from glycolysis and FAAO towards glucose oxidation. This confirms that LuCEMe can be used to detect metabolic changes in cells of interest in multicellular culture systems.

## Results

### High glucose levels suppress mitochondrial dependence in cell line cultures

As the LuCEMe assay allows easy metabolic measurements on cells in culture, we first assessed the ability of our method to detect metabolic differences between cell types by profiling a panel of cancerous and non-cancerous cell lines from diverse species and tissue origins. This analysis revealed a consistent and unexpected high level of glucose-supported ATP production and comparatively limited mitochondrial dependence in most cell lines tested. The glycolytic capacity and glucose dependency were uniformly high, whereas the mitochondrial dependence and FAAO capacity associated with mitochondrial ATP generation remained modest across conditions (Figure 2A). This suggests that, under in vitro conditions, metabolic phenotypes converge toward a predominantly glycolytic state, irrespective of genetic background, cell state or tissue of origin.

**Figure 2:**
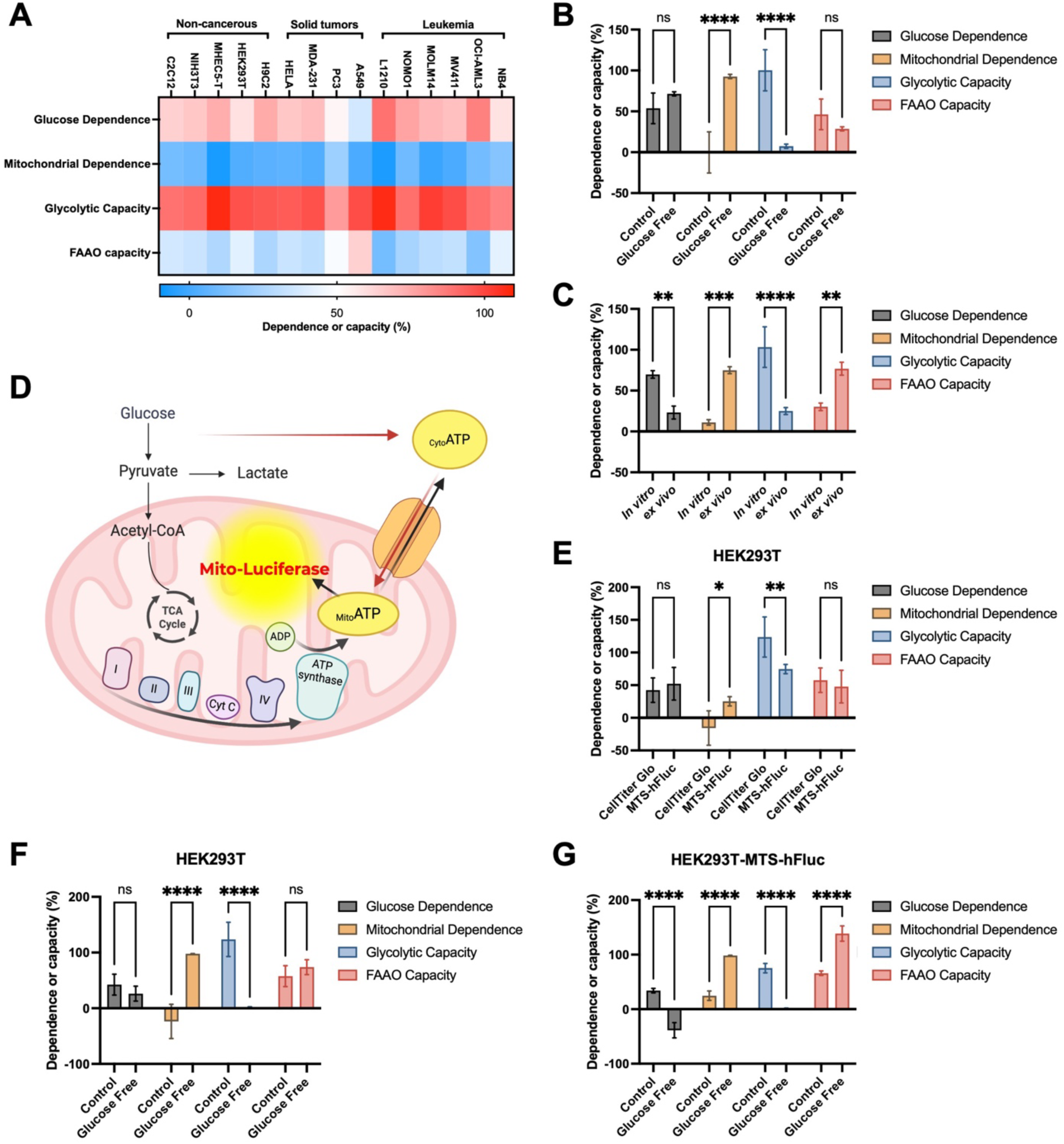
High glucose levels suppress mitochondrial dependence in cell line cultures. **(A)** Metabolic profile of a variety of different in vitro-cultured cell lines, including leukemia cells, solid tumor cells and non-cancerous cells, after LuCEMe analysis **(B)** The metabolic profile of MLL-AF9 AML cells after culture in glucose-free media for 24 hours, after LuCEMe analysis. **(C)** The metabolic profile of freshly isolated MLL-AF9 AML cells from mice in comparison with in vitro cultured AML cells, revealed by LuCEMe analysis. **(D)** Schematic showing the concept of the mitochondrial luciferase modification for LuCEMe analysis. **(E)** Comparing the metabolic profile of HEK293T cells after measurement of either total ATP (CellTiter Glo) or mitochondrial ATP (MTS-hFluc). **(F-G)** The metabolic profile of HEK293T after measurement of either total ATP (F) or mitochondrial ATP (G) in glucose-free culture conditions. Data are represented as mean ± SD. *p < 0.05, **p < 0.01, ***p < 0.001.

To exclude the possibility that the lack of detection of mitochondrial dependence for ATP synthesis across cell lines was due to an artefact of our method, we cultured our AML mouse cells in glucose free media for 24 hours and observed a strong shift towards mitochondrial ATP production with a complete lack of glycolytic capacity (Figure 2B). Furthermore, we compared the metabolic profiles of MLL-AF9 AML cells maintained in vitro with those of the same cells following engraftment in recipient mice and subsequent ex vivo isolation. AML cells recovered from the bone marrow displayed a marked shift in their energetic profile, characterized by increased mitochondrial dependence accompanied by reduced glucose dependence relative to their in vitro counterparts (Figure 2C). Together, these observations are consistent with the notion that standard culture conditions impose a metabolic phenotype favoring glucose utilization and highlight the ability of our method to detect such metabolic convergences.

Since the LuCEMe method measures metabolic dependencies and capacities, it is possible that baseline mitochondrial activity of cell lines is overlooked due to rapid metabolic switching in cultures characterized by glucose overabundance. To assess this possibility and test whether mitochondrial contributions to ATP production could be detected under basal culture conditions using LuCEMe, we applied a modified version of our method by expressing a mitochondria-targeted luciferase in HEK293T cells, which should enable the selective quantification of mitochondrial ATP independent of glycolytic contributions (Figure 2D). In comparison to the global ATP measurements, direct measurement of mitochondrial ATP revealed a modest mitochondrial dependence in HEK293T cells under normal culture conditions (Figure 2E), as revealed by a significant decrease of ATP levels upon OG treatment (Supplemental Figure 2A). However, in comparison to total ATP levels, the fraction attributable to mitochondrial pathways remained limited, while glucose dependence for mitochondrial ATP remained the same as for total ATP levels (Figure 2E). Glycolytic capacity (i.e. the fraction of ATP levels that can be maintained when OXPHOS is blocked) measured using the mitochondria-targeted luciferase was lower than when measured using CellTiter Glo but was still quite substantial, indicating either that ATP generated through glycolysis in the cytosol is shuttled into the mitochondria in OG-treated HEK293T cells ^13,14^, or that the mitochondria-targeted luciferase is not purely specific for mitochondrial ATP levels.

To further evaluate the extent to which glucose metabolism can underemphasize baseline mitochondrial parameters, we compared the measurement of total ATP and mitochondrial ATP in HEK293T cells cultured under glucose-free conditions. Similarly to our AML mouse cells, total ATP measurements in the absence of glucose revealed a pronounced loss of the glycolytic capacity and a corresponding increase in mitochondrial dependence for ATP production as expected (Figure 2F). Despite this shift toward mitochondrial metabolism, no substantial changes were observed in the FAAO capacity when inferred from total ATP measurements. Furthermore, glucose dependence remains unchanged as 2DG can still induce a significant decrease of total ATP even in glucose-free conditions (Supplemental Figure 2B). This suggests that compensatory mechanisms such as usage of alternative sugars like fructose or mobilization of glycogen stores can sustain low levels of glycolysis. Measurement of mitochondrial ATP levels in glucose-free conditions did show a strong reduction of glucose dependence and glycolytic capacity and a compensatory increase in mitochondrial dependence and FAAO capacity (Figure 2G). These results show that, while total ATP measurements capture global shifts in pathway dependence, direct mitochondrial ATP quantification provides substrate-level resolution of oxidative metabolism that can otherwise be masked by compensatory mechanisms.

### Adaptation of LuCEMe assay allows mitochondrial activity measurement

While using a mitochondria-targeting luciferase in the LuCEMe assay can give more insights into mitochondrial metabolism, this strategy requires genetic modification of target cells, making it less amenable to routine laboratory practice. To enable resolution of mitochondrial activity in any cell, we implemented another modification to the assay, designed to uncouple glycolytic inhibition from suppression of glucose-supported mitochondrial oxidation. 2DG treatment effectively suppresses not only glycolytic ATP production but also mitochondrial ATP generation from glucose-derived pyruvate oxidation, thus masking the contribution of mitochondria to total ATP production under standard culture conditions. By supplementing exogenous pyruvate during the assay, this upstream blockade of pyruvate oxidation imposed by 2DG can be bypassed, enabling therefore assessment of mitochondrial activity while maintaining standard culture glucose levels (Figure 3A).

**Figure 3:**
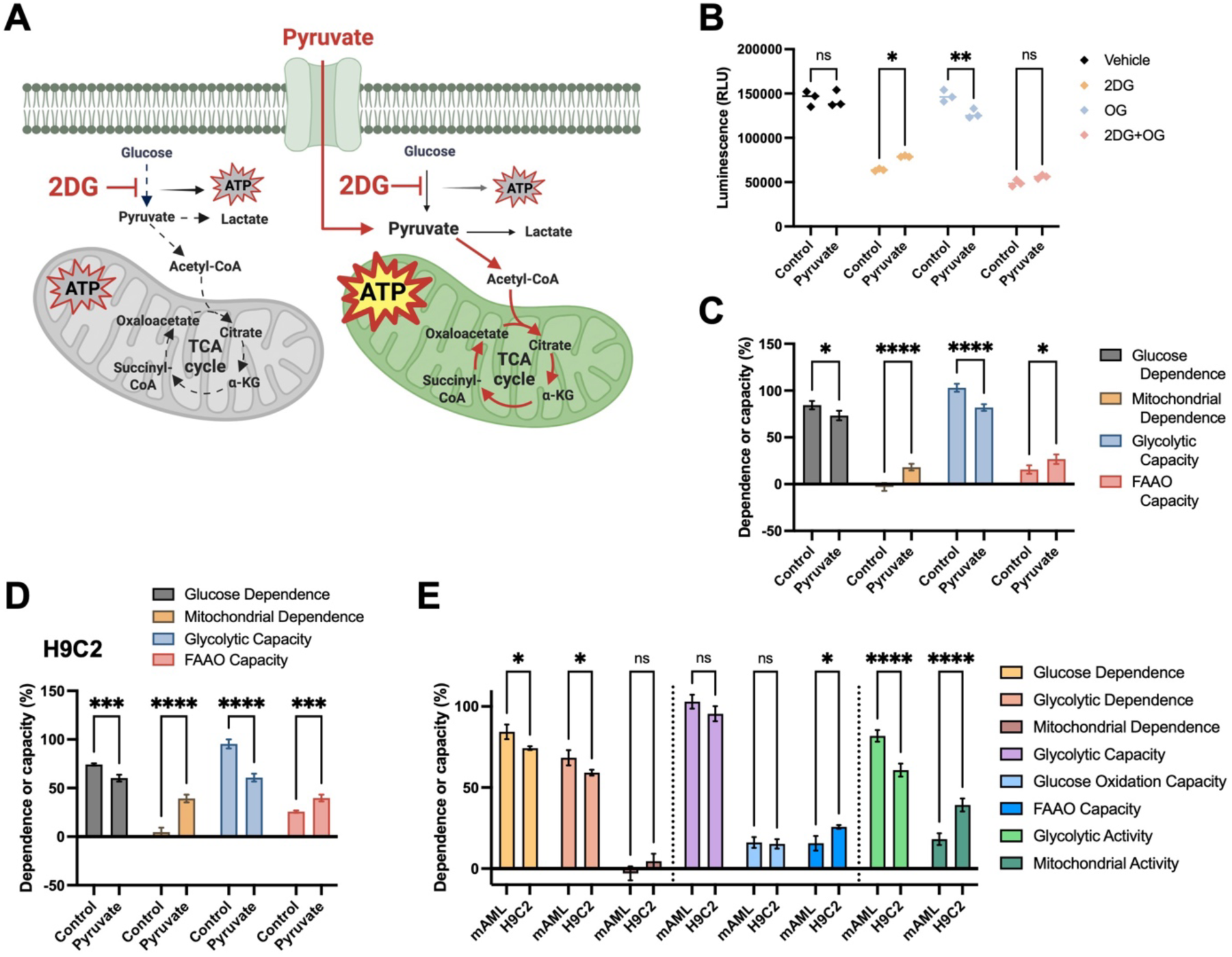
Adaptations of LuCEMe assay allows mitochondrial activity measurement. **(A)** LuCEMe modification concept to uncouple the effect of 2DG on glycolysis and glucose oxidation using exogenous pyruvate. **(B)** Total ATP levels of pyruvate treated MLL-AF9 AML cells after exposure to 2DG, OG and the combination. **(C-D)** The metabolic profile of MLL-AF9 and H9C2 respectively using the pyruvate LuCEMe modification. **(E)** Comparison of the metabolic activities of MLL_AF9 AML and H9C2 cells using the pyruvate LuCEMe modification. Data are represented as mean ± SD. *p < 0.05, **p < 0.01, ***p < 0.001.

Using this approach on our mouse luciferase-expressing AML cells, we detected a significant reduction in total ATP levels upon OG exposure while the 2DG-induced decrease of ATP levels was less pronounced in the presence of pyruvate, enabling the detection of a low baseline mitochondrial dependence (Figure 3B, 3C). Given that AML cells in culture are known to generate ATP mostly from glycolysis^15^, we also tested whether this modified LuCEMe assay could reveal the baseline mitochondrial activity of H9C2 cells, a rat cardiomyoblast cell line known to have a more mitochondria-dependent metabolism^16^. Under standard assay conditions, this mitochondrial activity was not readily detectable, likely due to the dominant contribution of glucose metabolism to both glycolysis and OXPHOS and effects of 2DG on both (Figure 2A). In contrast, pyruvate supplementation revealed a strong mitochondrial contribution to baseline ATP production in H9C2 cells (Figure 3D).

Importantly, this strategy also enabled the calculation of additional metabolic parameters in intact cells without the need for mitochondrial luciferase expression or glucose withdrawal. Mitochondrial activity, glycolytic activity, glycolytic dependence as well as glucose oxidation capacity can be inferred from ATP measurements obtained following treatment with 2DG, OG, and their combination in the presence or absence of pyruvate. Using this framework, we identified clear metabolic distinctions between cell types, with H9C2 cells exhibiting higher mitochondrial activity predominantly supported by FAAO, while mouse MLL-AF9 AML cells showed stronger glycolytic activity in combination with modest mitochondrial activity supported by both glucose oxidation and FAAO (Figure 3E).

### LuCEMe assay reveals metabolic changes during cell differentiation and coculture

Having established the sensitivity and specificity of LuCEMe through a series of technical validations and metabolic perturbations, we next sought to determine whether this approach could capture biologically relevant metabolic adaptations in more complex cellular contexts. To this end, we applied the method to models of cellular differentiation and to coculture systems comprising distinct cell populations.

We first analyzed metabolic changes occurring during macrophage polarization, which was induced using either a combination of lipopolysaccharide (LPS) and interferon gamma (IFNγ) to promote a pro-inflammatory phenotype characterized by increased expression of CD80 or by adding IL-4 to drive an anti-inflammatory state characterized by increased CD206 expression (Supplemental Figure 3) ^17^. Metabolic profiling using LuCEMe was performed immediately after polarization and revealed a shift from mitochondrial dependence toward increased glycolytic capacity upon polarization, while glucose dependence and FAAO capacity remained largely unchanged (Figure 4A). Interestingly, the metabolic changes were nearly identical between macrophages polarized towards a pro- or anti-inflammatory state. This is different from results obtained with SCENITH or other protein synthesis-based metabolic assays, which revealed almost no changes in mitochondrial dependence or glycolytic capacity in anti-inflammatory macrophages^18^, but is in line with a previous report showing enhanced glycolytic engagement in activated macrophages regardless of polarization type as revealed by metabolomics analysis^19^.

**Figure 4:**
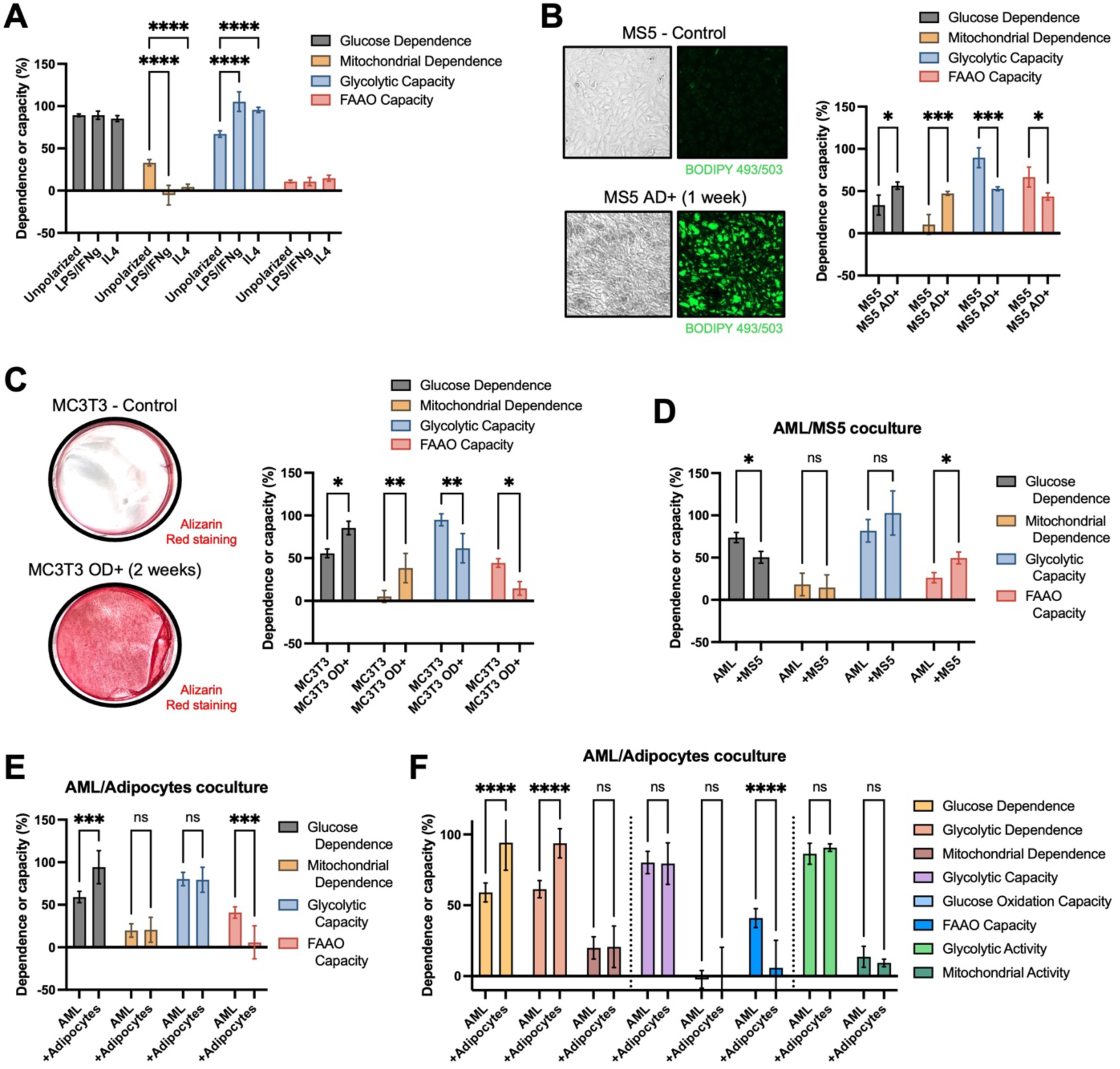
LuCEMe reveals metabolic changes during cell differentiation and coculture. **(A)** The metabolic profile of in vitro cultured mouse macrophages after polarization for 24 hours with either LPS/IFNγ or IL4 using LuCEMe analysis. **(B)** LuCEMe analysis of MS5 stromal cells before and after adipogenic differentiation for 1 week, prior to metabolic profiling. Images reveal lipid droplets stained using BODIPY 493/503. **(C)** LuCEMe analysis of MC3T3 osteoprogenitor cells before and after osteogenic differentiation for 2 weeks, prior to metabolic profiling. Images show alizarin red staining to reveal mineral deposits. **(D)** The metabolic profile of MLL-AF9 AML cells after coculture with undifferentiated MS5 cells for 24 hours. **(E)**The metabolic profile of MLL-AF9 AML cells after coculture with adipogenically-differentiated MS5 cells for 24 hours. **(F)** LuCEMe assay with pyruvate modification showing the metabolic activity of MLL-AF9 AML cells after coculture with adipogenically-differentiated MS5 cells. Data are represented as mean ± SD. *p < 0.05, **p < 0.01, ***p < 0.001.

Next, we examined metabolic changes during differentiation using MS5 mouse bone marrow-derived mesenchymal stromal cells. Adipogenic differentiation induced a measurable shift in metabolic state, characterized by increased glucose and mitochondrial dependence accompanied by a reduction in glycolytic and FAAO capacity (Figure 4B). This profile indicates a constrained yet highly engaged metabolic state, consistent with the elevated energetic and biosynthetic demands associated with lipid synthesis and storage (Figure 4B). We also assessed metabolic remodeling during osteogenic differentiation using the mouse MC3T3 osteoprogenitor cell line. Like the adipogenic differentiation, our LuCEMe assay revealed an increased dependence on both glucose and mitochondrial ATP production, while an overall reduction in metabolic capacity was observed, with both glycolytic and FAAO capacities diminished following differentiation (Figure 4C). This shift is consistent with the elevated energetic demands associated with matrix production and mineralization in osteoblasts (Figure 4C). Together, these findings show a transition toward a more constrained but highly active metabolic state during adipogenic or osteogenic differentiation of mesenchymal stromal cells, in line with the current consensus^20^.

The LuCEMe assay allows to investigate cellular metabolism in cocultures by cell type-specific luciferase expression, as we showed in AML cell-macrophage cocultures (Figure 1F, 1G). To further investigate metabolic adaptations arising from cell-cell interactions within a shared microenvironment, we performed LuCEMe on AML cells after coculture with MS5 mesenchymal stromal cells or differentiated mature adipocytes. Interestingly, the metabolic changes occurring in AML cells cocultured with stromal cells or adipocytes were quite distinct from those observed in cocultures with macrophages. AML cells cocultured with MS5 cells had higher ATP levels compared to monocultures, like observed in macrophage cocultures, but 2DG alone had less effects on ATP levels, while the combination of 2DG with OG led to a more pronounced decrease of ATP in cocultures (Supplemental Figure 4A). Together, this revealed a decreased glucose dependence and increased FAAO capacity in AML cells after coculture with MS5 cells (Figure 4D), while AML cells in coculture with macrophages showed increased glucose dependence linked to higher mitochondrial glucose oxidation (Figure 1G). These findings are in line with numerous studies that have demonstrated how stromal interactions fundamentally reshape leukemia cell bioenergetics^21^ and confirm a previous report showing that mitochondrial OXPHOS activity is significantly elevated when AML cells are cocultured with mesenchymal stromal cells^22^. Coculturing AML cells with adipocytes revealed yet another metabolic profile, with a strong increase in their glucose dependence, and a surprising collapse of their FAAO capacity (Figure 4E). Using the pyruvate modification of our LuCEMe assay we confirmed an increased glycolytic dependence and strongly reduced FAAO capacity in AML cells co-cultured with adipocytes, but no significant changes to baseline glycolytic and mitochondrial activity were found (Figure 4F), suggesting that competition for exogenous fatty acids renders AML cells almost fully dependent on glycolysis for ATP production.

To assess whether coculture-induced metabolic changes were unique to AML cells, we performed LuCEMe on the bone-metastatic breast cancer cell line MDA-MB-231 after coculture with MS5 cells or the macrophage cell line. Coculturing MDA-MB-231 cells with MS5 cells revealed a similar metabolic phenotype to AML cells, characterized by reduced glucose dependency and increased FAAO capacity (Supplemental Figure 4B), while surprisingly no significant differences were observed after coculture with macrophages, whether these were unpolarized or polarized towards a pro-inflammatory phenotype (Supplemental Figure 4C).

Taken together, these findings highlight the strength of the LuCEMe assay, revealing that distinct cancer types undergo unique metabolic adaptations when interacting with different cells of the tumor microenvironment.

### Ex vivo LuCEMe analysis reveals new insights into AML and T cell metabolism

We next evaluated whether LuCEMe could capture the metabolic profiles of cells in their native environment by performing rapid ex vivo analysis. As a first proof-of-principle, we assessed the metabolic profile of ^CD3+CD8+^T cells after activation, known to increase their glycolytic activity. To this end, mice were primed with LPS and rechallenged one week later with a higher dose to elicit an adaptive immune response^23^. ^CD3+CD8+^T cells were isolated 24 hours following rechallenge, and activation was confirmed by upregulation of canonical markers, including Ly6C, CD44, and CD25 (Figure 5A). Metabolic profiling of freshly isolated ^CD3+CD8+^T cells at baseline revealed a state characterized by minimal glycolytic capacity, modest FAAO capacity, and a high dependence on both mitochondrial and glucose-derived ATP production, indicating glucose oxidation as the primary metabolic pathway active in these cells. Following LPS rechallenge, ^CD3+CD8+^T cells exhibited a metabolic shift marked by a modest increase in glucose dependence and significantly enhanced glycolytic capacity, accompanied by a reduction in mitochondrial dependency and FAAO capacity (Figure 5B). Thus, our LuCEMe-based metabolic analysis reflects an increase in glycolysis in activated ^CD3+CD8+^T cells, consistent with the current consensus in the field^24^.

**Figure 5:**
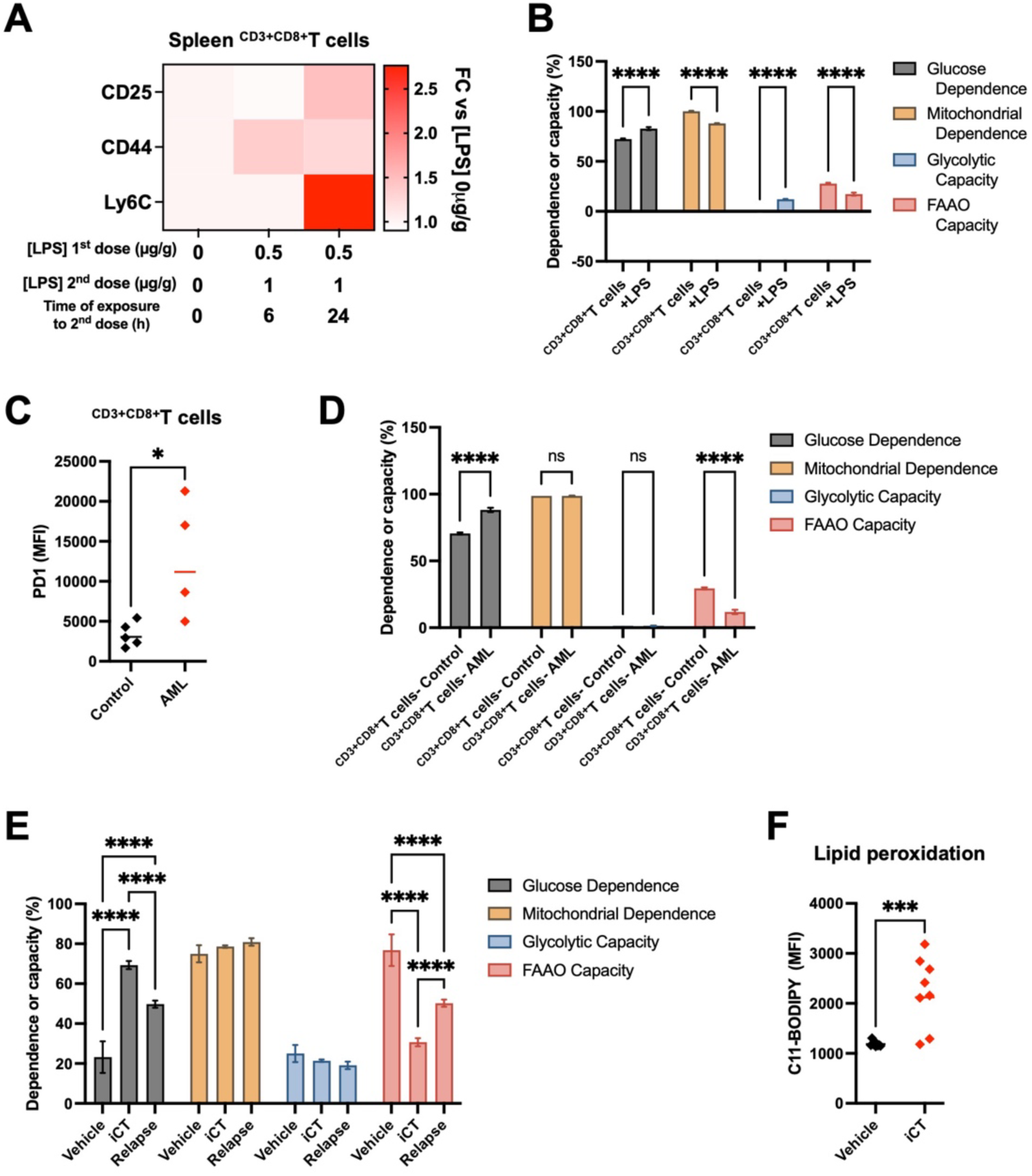
Ex vivo metabolic profiling using LuCEMe reveals new insights into AML and T cell metabolism. **(A)** Heatmap representing the expression levels of T cell activation markers in splenic ^CD3+CD8+^T cells of mice that were rechallenged with LPS for either 6 or 24 hours prior to analysis by flow cytometry. FC: Fold change. **(B)** The metabolic profile of CD8^+^ T cells in LPS rechallenged mice, at 24 hours after the second dose, revealed by ex vivo LuCEMe analysis. **(C)** PD1 expression in ^CD3+CD8+^T cells from leukemic mice. MFI: Mean fluorescence intensity. **(D)** The metabolic profile of ^CD3+CD8+^T cells in leukemic mice revealed by ex vivo LuCEMe analysis. **(E)** The metabolic profile of MLL-AF9 AML cells obtained from mice with vehicle treatment, at 3 days after induction chemotherapy (iCT; 5 days of cytarabine 100 mg/kg + 3 days of doxorubicin 3 mg/kg, i.p.) or at 10 days after iCT (relapse), revealed by ex vivo LuCEMe analysis. **(F)** Levels of lipid peroxidation measured using C11-BODIPY staining in MLL-AF9 AML cells obtained from mice before or 3 days after iCT. Data are represented as mean ± SD.*p < 0.05, **p < 0.01, ***p < 0.001.

We next assessed whether LuCEMe could resolve metabolic alterations associated with T cell dysfunction in a disease context. To this end, ^CD3+CD8+^T cells were isolated from mice bearing AML or from control animals, and metabolic profiling was performed immediately ex vivo. Consistent with the establishment of an exhausted phenotype, ^CD3+CD8+^T cells from leukemic mice exhibited increased expression of the inhibitory receptor PD1 (Figure 5C). Metabolic profiling with LuCEMe revealed an increased dependence on glucose alongside a strong reduction in FAAO capacity. Notably, this shift occurred without significant changes in glycolytic capacity or overall mitochondrial dependence (Figure 5D), suggesting a failure to switch to a glycolytic profile and a shift towards even higher glucose oxidation. While exhausted ^CD3+CD8+^T cells generally exhibit metabolic dysfunction characterized by a decreased glycolytic capacity and decreased OXPHOS activity^25^, prior analysis in AML models indicated a similar impaired T cell metabolic switching characterized by an abnormally elevated OXPHOS-high metabolic state^26^.

Finally, we sought to determine whether LuCEMe could uncover metabolic adaptations in AML cells during disease progression and treatment response. Our own previous studies and others have shown, using transcriptomic analyses and extracellular flux measurements, that AML cells persisting through chemotherapy exhibit increased OXPHOS activity and enhanced fatty acid and glutamine metabolism^11,12^. However, these approaches do not directly resolve substrate-specific contributions to ATP production, and it remains unclear if residual AML cells exhibit increased fatty acid or amino acid oxidation. To address this question, we isolated AML cells isolated from mice prior to chemotherapy, at the time of maximal therapeutic response, and at relapse, followed by immediate ex vivo metabolic profiling using LuCEMe. Using this approach, we identified a distinct and previously unrecognized metabolic shift occurring at the time of maximal response to chemotherapy. Specifically, AML cells persisting through chemotherapy exhibited a significant increase in glucose dependence accompanied by a marked reduction in FAAO capacity (Figure 5E). Notably, this shift occurred without detectable changes in overall mitochondrial dependence or glycolytic capacity, indicating that the total contribution of these pathways to ATP production remains but that the cells shift towards oxidation of glucose rather than fatty acids or amino acids. This phenotype persisted through relapse, suggesting a durable rewiring of metabolic substrate utilization.

The significantly decreased FAAO capacity in residual AML cells was unexpected, as an increased fatty acid metabolism signature has been consistently reported in AML cells persisting after chemotherapy in both mouse models and patient samples^11,12^. Given that residual AML cells also exhibit high ROS levels^27^ reflecting their increased OXPHOS activity, one possibility is that changes in fatty acid metabolism serve to counteract excessive lipid peroxidation. In line with this, we found that lipid peroxidation is increased in AML cells at the time of maximal response to chemotherapy in vivo (Figure 5F).

These findings indicate that, rather than a fatty acid-fueled increase in oxidative metabolism, AML cells undergoing therapeutic stress preferentially redirect glucose towards mitochondrial oxidation for ATP production, thus liberating fatty acid pools to counteract lipid peroxidation. This observation reveals a previously unrecognized metabolic state associated with treatment response, highlighting the ability of LuCEMe to resolve adaptations that are not captured by conventional approaches.

## Discussion

A central challenge in the study of cellular metabolism is the ability to directly assess metabolic pathway activity within native, unperturbed environments while preserving physiologically relevant cell-cell interactions. Most current approaches require either substantial manipulation of culture conditions or rely on indirect readouts that can obscure the functional output of metabolism. Here, we present LuCEMe, a framework that enables direct measurement of metabolic pathways sustaining cellular ATP production under standard culture conditions, while remaining fully compatible with coculture systems. This capability allows basic metabolic phenotyping to be performed without disrupting the native cellular context, thereby capturing both intrinsic metabolic states and adaptations driven by intercellular interactions. By combining selective perturbations of metabolic pathways with direct ATP measurements, our approach resolves the relative contribution of glycolytic and mitochondrial pathways, as well as the substrates fueling mitochondrial ATP, providing a quick, reliable and affordable strategy to interrogate metabolism across in vitro, ex vivo, and multicellular settings. By measuring ATP production directly and resolving pathway level contributions in native culture conditions without the need for cell separation prior to analysis, LuCEMe complements existing techniques while addressing a key gap in functional metabolic analysis.

Across the diverse systems examined in the current study, several unifying principles emerge. Firstly, glucose availability exerts a masking effect on mitochondrial metabolism. Under standard culture conditions, a wide range of cell types converge toward a glucose-dependent phenotype with little necessity for mitochondrial ATP production, obscuring underlying differences in basal mitochondrial activity. By directly measuring mitochondrial ATP production or by supplementing exogenous pyruvate to uncouple pyruvate oxidation from glycolysis, heterogeneity in mitochondrial ATP production activity and capacity can be revealed that is otherwise not apparent.

Secondly, leveraging the ability of LuCEMe to study cellular metabolism in intact cocultures, we found that cancer cells undergo specific metabolic adaptations depending on which other cell type they are exposed to, and that these adaptations are cancer-type dependent. Mouse MLL-AF9 AML cells for example show massive activation of glucose oxidation when cocultured with macrophages but shift to FAAO when in the presence of mesenchymal stromal cells or become highly dependent on glycolysis when competing with adipocytes. Human MDA-BM-231 breast cancer cells in contrast do not change their metabolism in the presence of macrophages, but do increase FAAO in coculture with mesenchymal stromal cells like the AML cells. Thirdly, our LuCEMe analyses revealed that metabolic remodeling in response to differentiation, activation, or stress is often characterized not by a simple shift between glycolysis and OXPHOS, but by a more constrained metabolic state in which overall capacity is reduced while dependence on specific pathways increases. This pattern was consistently observed across differentiation models, immune activation states, and disease contexts.

One of the most notable insights enabled by LuCEMe arises from the analysis of AML cells during chemotherapy treatment. While prior studies have associated chemoresistance with increased OXPHOS and enhanced fatty acid metabolism^11,12^, these conclusions were mostly based on transcriptomic signatures or proxy measurements of metabolic activity. By directly quantifying ATP production, we identify a distinct metabolic state at the time of maximal response to chemotherapy, characterized by increased glucose dependence and reduced FAAO capacity, without changes in overall mitochondrial dependence or glycolytic capacity. This indicates a selective rewiring of substrate utilization for mitochondrial ATP production, rather than a global shift in metabolic pathway engagement. The partial persistence of this phenotype through relapse suggests that therapeutic pressure may induce durable changes in metabolic wiring that are not captured by conventional approaches. Interestingly, the discovery that AML cells persisting after chemotherapy may redirect fatty acids from mitochondrial oxidation towards the remediation of lipid peroxidation provides an explanation for the previous observation that AML cells lacking ALDH3A2, an enzyme responsible for detoxification of fatty aldehydes resulting from lipid peroxidation, are much more sensitive to chemotherapy^28^. More broadly, these findings illustrate how resolving ATP production at the substrate level can reveal biologically and clinically relevant metabolic states that would otherwise remain obscured.

Looking forward, the LuCEMe framework opens new opportunities for the study of metabolism in physiologically relevant contexts. Its compatibility with low cell numbers, primary cells and ex vivo assays enables metabolic profiling without extensive manipulation, which helps preserve the native metabolic state of the cells. Its applicability to coculture systems allows interrogation of microenvironment-driven metabolic interactions. In particular, the ability to resolve pathway-specific ATP production in multicellular settings provides a means to dissect how stromal and immune components shape cellular metabolism in health and disease. In therapeutic contexts, identifying shifts in substrate utilization may reveal metabolic vulnerabilities that are not apparent from extracellular flux and metabolomics analyses alone. More broadly, by directly linking metabolic pathways to their functional output, this approach provides a foundation for a more precise and integrative understanding of cellular metabolism.

### Limitations of the study

Despite the strengths of the LuCEMe assay, some limitations should be considered. First, while ATP production represents a central functional output of cellular metabolism, it does not capture the full complexity of metabolic fluxes supporting biosynthesis or redox regulation. LuCEMe may thus represent the ideal method to quickly and easily perform a first assessment of cellular metabolism, followed by more detailed metabolic profiling through metabolomics analysis, stable isotope tracing and/or metabolic enzyme perturbations. Second, the interpretation of pathway and substrate contributions relies on the specificity and efficacy of pharmacological perturbations, which may vary across cell types and experimental conditions.

In the context of coculture systems, an additional layer of complexity arises as it cannot be formally excluded that metabolic phenotypes observed in one cell population reflect indirect effects mediated by pharmacological perturbations acting on a second cell type. For example, inhibition of a metabolic pathway in one cell population may alter substrate availability or secreted factors, thereby inducing secondary metabolic adaptations in neighboring cells. As such, distinguishing direct cell-intrinsic responses from microenvironment-driven effects remains a challenge when relying solely on pharmacological interventions. A potential strategy to address this limitation is the implementation of genetically encoded, inducible perturbation systems targeting key rate-limiting enzymes of glycolytic and mitochondrial ATP producing pathways. This would enable cell-type-specific modulation of metabolic pathways while preserving the multicellular context, allowing more precise attribution of ATP production changes to intrinsic versus extrinsic mechanisms. Coupling such systems with LuCEMe would provide a powerful framework to dissect metabolic interactions within complex cellular environments.

## Acknowledgments

We thank Nicolas Dauguet from the CYTF platform at the de Duve Institute for assistance with cell sorting. We are also grateful to Justine van de Velve and Pierre Sonveaux (IREC, UCLouvain, Belgium) for providing the MDA-MB-231 cell line expressing firefly luciferase. Figures 1A, 2D and 3A were created with BioRender.com. This work was supported by grants from the Belgian Foundation Against Cancer (F/2020/1440, F/2024/2556), the Fund for Scientific Research - FNRS (F.R.S.-FNRS; A5/5-CQ/135, M4/1/2/5-MIS/BEJ), the Special Funds for Research (FSR; Fédération Wallonie-Bruxelles - UCLouvain) and the de Duve Institute. L.L. was supported by a doctoral research fellowship from the Fund for Research Training in Industry and Agriculture (FRIA; FRIA/FC-4456] and by a Bourse du Patrimoine from the UCLouvain. C.S. was supported by a Télévie Grant from the F.R.S.-FNRS (F5/20/5-TLV/BC).

## Author Contributions

L.L. and N.v.G. conceptualized the study and designed experiments. L.L. and C.S. performed experiments. L.L., C.S. and N.v.G. analyzed and interpreted results. L.L. and N.v.G. wrote the manuscript text and prepared the figures. N.v.G. acquired funding and supervised the research. All authors reviewed the manuscript.

## STAR Methods

### KEY RESOURCES TABLE

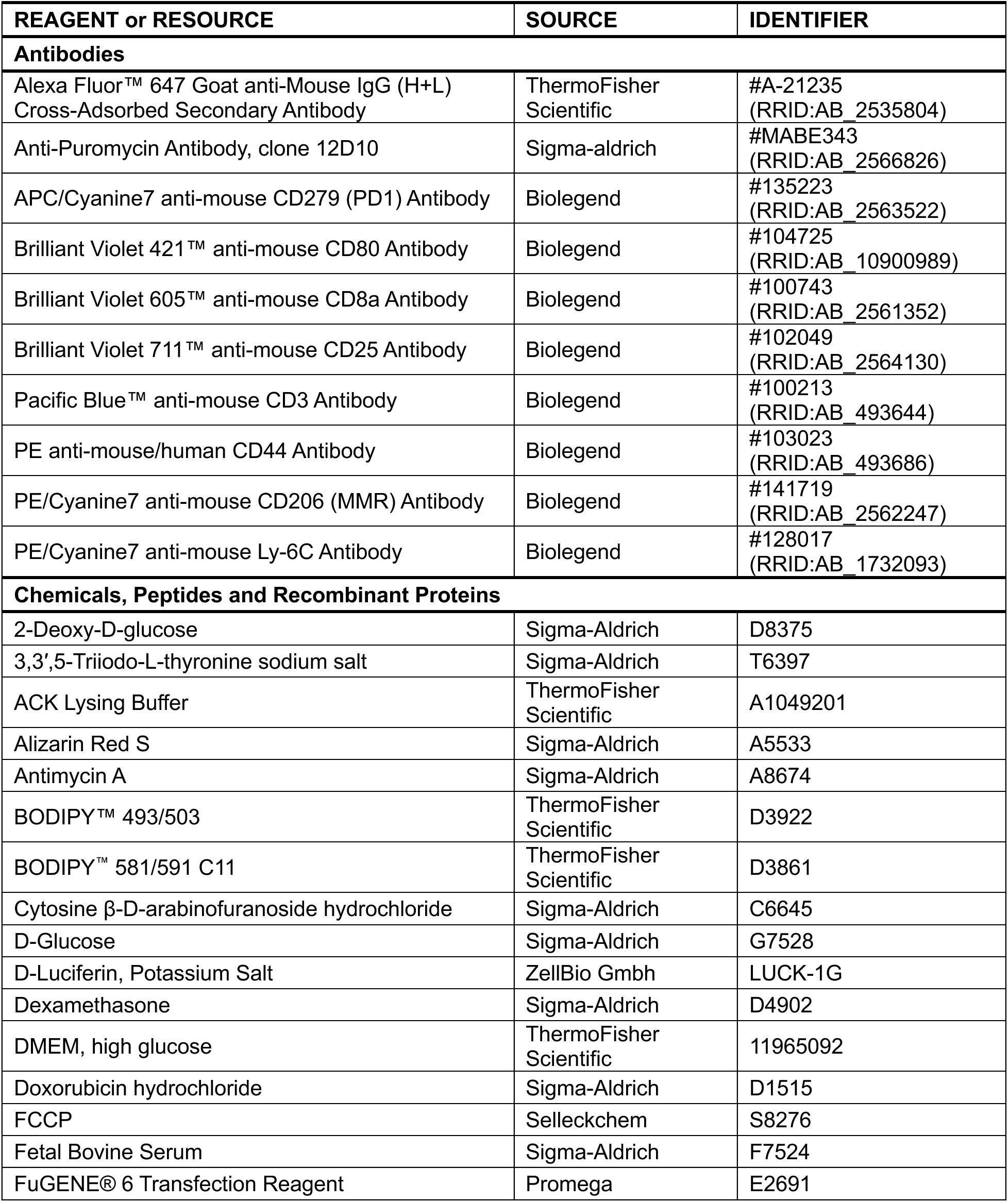
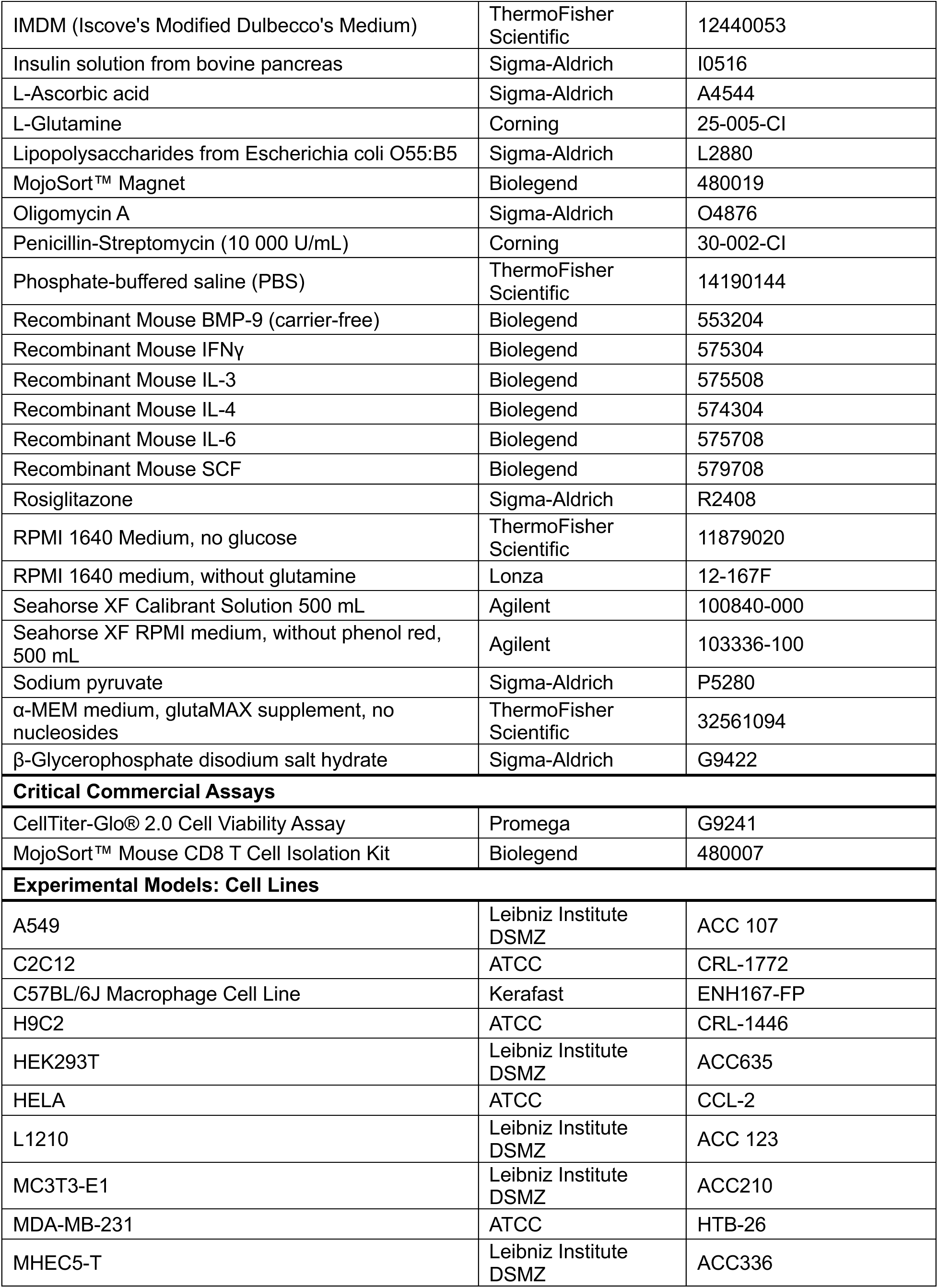
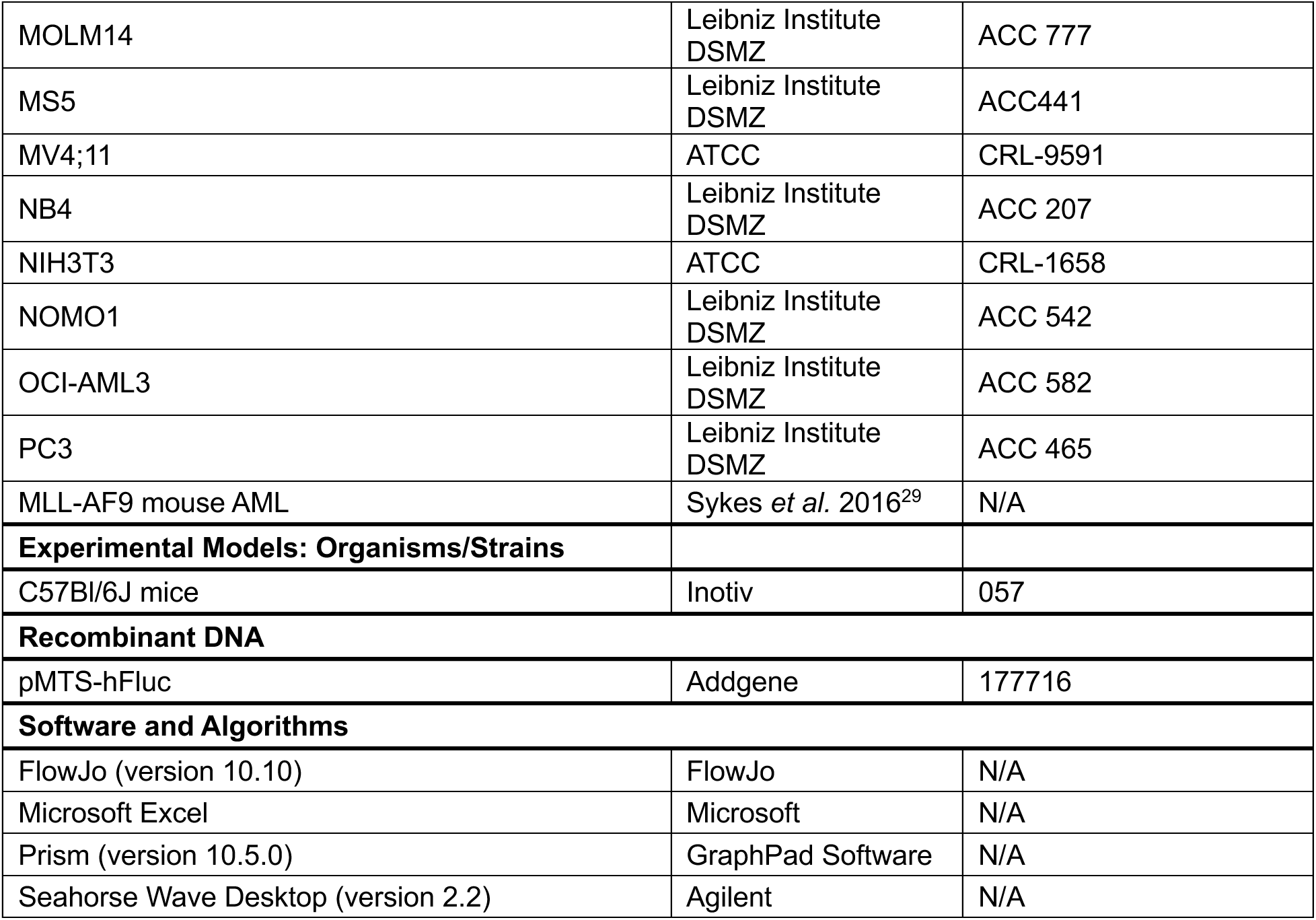

| REAGENT or RESOURCE | SOURCE | IDENTIFIER |
|---|---|---|
| **Antibodies** |  |  |
| Alexa Fluor™ 647 Goat anti-Mouse IgG (H+L) Cross-Adsorbed Secondary Antibody | ThermoFisher Scientific | #A-21235 (RRID:AB_2535804) |
| Anti-Puromycin Antibody, clone 12D10 | Sigma-aldrich | #MABE343 (RRID:AB_2566826) |
| APC/Cyanine7 anti-mouse CD279 (PD1) Antibody | Biolegend | #135223 (RRID:AB_2563522) |
| Brilliant Violet 421™ anti-mouse CD80 Antibody | Biolegend | #104725 (RRID:AB_10900989) |
| Brilliant Violet 605™ anti-mouse CD8a Antibody | Biolegend | #100743 (RRID:AB_2561352) |
| Brilliant Violet 711™ anti-mouse CD25 Antibody | Biolegend | #102049 (RRID:AB_2564130) |
| Pacific Blue™ anti-mouse CD3 Antibody | Biolegend | #100213 (RRID:AB_493644) |
| PE anti-mouse/human CD44 Antibody | Biolegend | #103023 (RRID:AB_493686) |
| PE/Cyanine7 anti-mouse CD206 (MMR) Antibody | Biolegend | #141719 (RRID:AB_2562247) |
| PE/Cyanine7 anti-mouse Ly-6C Antibody | Biolegend | #128017 (RRID:AB_1732093) |
| **Chemicals, Peptides and Recombinant Proteins** |  |  |
| 2-Deoxy-D-glucose | Sigma-Aldrich | D8375 |
| 3,3′,5-Triiodo-L-thyronine sodium salt | Sigma-Aldrich | T6397 |
| ACK Lysing Buffer | ThermoFisher Scientific | A1049201 |
| Alizarin Red S | Sigma-Aldrich | A5533 |
| Antimycin A | Sigma-Aldrich | A8674 |
| BODIPY™ 493/503 | ThermoFisher Scientific | D3922 |
| BODIPY™ 581/591 C11 | ThermoFisher Scientific | D3861 |
| Cytosine β-D-arabinofuranoside hydrochloride | Sigma-Aldrich | C6645 |
| D-Glucose | Sigma-Aldrich | G7528 |
| D-Luciferin, Potassium Salt | ZellBio Gmbh | LUCK-1G |
| Dexamethasone | Sigma-Aldrich | D4902 |
| DMEM, high glucose | ThermoFisher Scientific | 11965092 |
| Doxorubicin hydrochloride | Sigma-Aldrich | D1515 |
| FCCP | Selleckchem | S8276 |
| Fetal Bovine Serum | Sigma-Aldrich | F7524 |
| FuGENE® 6 Transfection Reagent | Promega | E2691 |
| IMDM (Iscove's Modified Dulbecco's Medium) | ThermoFisher Scientific | 12440053 |
| Insulin solution from bovine pancreas | Sigma-Aldrich | I0516 |
| L-Ascorbic acid | Sigma-Aldrich | A4544 |
| L-Glutamine | Corning | 25-005-CI |
| Lipopolysaccharides from Escherichia coli O55:B5 | Sigma-Aldrich | L2880 |
| MojoSort™ Magnet | Biolegend | 480019 |
| Oligomycin A | Sigma-Aldrich | O4876 |
| Penicillin-Streptomycin (10 000 U/mL) | Corning | 30-002-CI |
| Phosphate-buffered saline (PBS) | ThermoFisher Scientific | 14190144 |
| Recombinant Mouse BMP-9 (carrier-free) | Biolegend | 553204 |
| Recombinant Mouse IFNγ | Biolegend | 575304 |
| Recombinant Mouse IL-3 | Biolegend | 575508 |
| Recombinant Mouse IL-4 | Biolegend | 574304 |
| Recombinant Mouse IL-6 | Biolegend | 575708 |
| Recombinant Mouse SCF | Biolegend | 579708 |
| Rosiglitazone | Sigma-Aldrich | R2408 |
| RPMI 1640 Medium, no glucose | ThermoFisher Scientific | 11879020 |
| RPMI 1640 medium, without glutamine | Lonza | 12-167F |
| Seahorse XF Calibrant Solution 500 mL | Agilent | 100840-000 |
| Seahorse XF RPMI medium, without phenol red, 500 mL | Agilent | 103336-100 |
| Sodium pyruvate | Sigma-Aldrich | P5280 |
| α-MEM medium, glutaMAX supplement, no nucleosides | ThermoFisher Scientific | 32561094 |
| β-Glycerophosphate disodium salt hydrate | Sigma-Aldrich | G9422 |
| **Critical Commercial Assays** |  |  |
| CellTiter-Glo® 2.0 Cell Viability Assay | Promega | G9241 |
| MojoSort™ Mouse CD8 T Cell Isolation Kit | Biolegend | 480007 |
| **Experimental Models: Cell Lines** |  |  |
| A549 | Leibniz Institute DSMZ | ACC 107 |
| C2C12 | ATCC | CRL-1772 |
| C57BL/6J Macrophage Cell Line | Kerafast | ENH167-FP |
| H9C2 | ATCC | CRL-1446 |
| HEK293T | Leibniz Institute DSMZ | ACC635 |
| HELA | ATCC | CCL-2 |
| L1210 | Leibniz Institute DSMZ | ACC 123 |
| MC3T3-E1 | Leibniz Institute DSMZ | ACC210 |
| MDA-MB-231 | ATCC | HTB-26 |
| MHEC5-T | Leibniz Institute DSMZ | ACC336 |
| MOLM14 | Leibniz Institute DSMZ | ACC 777 |
| MS5 | Leibniz Institute DSMZ | ACC441 |
| MV4;11 | ATCC | CRL-9591 |
| NB4 | Leibniz Institute DSMZ | ACC 207 |
| NIH3T3 | ATCC | CRL-1658 |
| NOMO1 | Leibniz Institute DSMZ | ACC 542 |
| OCI-AML3 | Leibniz Institute DSMZ | ACC 582 |
| PC3 | Leibniz Institute DSMZ | ACC 465 |
| MLL-AF9 mouse AML | Sykes *et al.* 2016<sup>29</sup> | N/A |
| **Experimental Models: Organisms/Strains** |  |  |
| C57Bl/6J mice | Inotiv | 057 |
| **Recombinant DNA** |  |  |
| pMTS-hFluc | Addgene | 177716 |
| **Software and Algorithms** |  |  |
| FlowJo (version 10.10) | FlowJo | N/A |
| Microsoft Excel | Microsoft | N/A |
| Prism (version 10.5.0) | GraphPad Software | N/A |
| Seahorse Wave Desktop (version 2.2) | Agilent | N/A |

## RESOURCE AVAILABILITY

### Lead Contact

Further information and requests for resources and reagents should be directed to and will be fulfilled by the Lead Contact, Nick van Gastel.

### Materials Availability

This study did not generate new unique materials.

### Data and Code Availability

All data supporting the findings of this study are available within the paper.

## EXPERIMENTAL MODEL AND SUBJECT DETAILS

### Cell Culture

All cell cultures were performed in an incubator at 37°C and 5% CO_2_. HELA cells were cultured in IMDM supplemented with 100 U/mL penicillin, 100 µg/mL streptomycin and 10% fetal bovine serum (FBS). MDA-MB-231, PC3 and the bone marrow macrophage cell line were cultured in DMEM supplemented with 2 mM L-glutamine, 100 U/mL penicillin, 100 µg/mL streptomycin and 10% FBS. The MLL-AF9 primary AML cells were expanded in RPMI1640 medium supplemented with 100 U/mL penicillin, 100 mg/mL streptomycin, 10% FBS, 2 mM L-glutamine, 20 ng/mL recombinant mouse Stem Cell Factor (rmSCF), 10 ng/mL recombinant mouse Interleukin-3 (rmIL-3) and 10 ng/mL rmIL-6. MS5, MC3T3, NIH3T3, C2C12 and H9C2 cell lines were cultured in α-MEM medium supplemented with 100 U/mL penicillin, 100 µg/mL streptomycin and 10% FBS. All other cell lines were cultured in RPMI1640 medium supplemented with 2 mM L-glutamine, 100 U/mL penicillin, 100 µg/mL streptomycin and 10% FBS.

### Cell differentiation and polarization

The bone marrow macrophage cell line was cultured at 50 000 cells/well in Thermo Scientific Nunc MicroWell white 96-well optical-bottom plates in RPMI1640 for 24 hours prior to the polarization. Then RPMI1640 supplemented with either a combination of 1 µg/mL LPS and 20 ng/mL rmIFNγ for pro-inflammatory polarization or 20 ng/mL rmIL-4 for anti-inflammatory polarization was added for 24 hours.

For osteogenic differentiation, MC3T3 cells were cultured at 15 000 cells/well in Thermo Scientific Nunc MicroWell white 96-well optical-bottom plates until full confluency was reached. Then osteogenic medium composed of α-MEM medium supplemented with 10 mM β-glycerophosphate, 50 µg/mL ascorbic acid and 100 ng/mL recombinant mouse Bone Morphogenetic Protein 9 (rmBMP-9) was added and refreshed twice a week. After 14 days, cultures were stained for mineral deposits using alizarin red staining to assess the differentiation state^30^.

For adipogenic differentiation, MS5 cells were cultured at 15 000 cells/well in Thermo Scientific Nunc MicroWell white 96-well optical-bottom plates until full confluency was reached. Then adipogenic medium composed of α-MEM medium supplemented with 1 ng/mL triiodothyronine, 100 nM rosiglitazone, 100 nM dexamethasone and 1 µg/mL insulin was added and refreshed twice a week. After 1 week, cultures were stained with BODIPY 493/503 according to the manufacturer’s instructions to reveal lipid droplets.

### Animal experiments and *in vivo* treatment

All animal procedures performed in this study were approved by the UCLouvain Institutional Animal Care and Use Committee (approval numbers 2021/UCL/MD/03 and 2024/UCL/MD/56). Experiments were conducted on female C57Bl/6 mice aged between 8 to 10 weeks. Recipient mice were injected intra-venously with 1 million AML cells (tail vein). Leukemia engraftment was monitored using bioluminescence imaging. For chemotherapy treatment, mice were treated with a combination of 100 mg/kg cytarabine once daily for 5 days and 3 mg/kg doxorubicin once daily for the first 3 days, delivered via intraperitoneal injection, starting at day 15 after engraftment. For ^CD3+CD8+^T cell activation experiments, mice were treated by intraperitoneal injection with a first dose of LPS at 0.5 µg/g body weight and then a second dose of 1 µg/g after one week. Mice were euthanized by CO_2_ asphyxiation for organ collection (bones and spleen) for further analyses.

## METHOD DETAILS

### Standard LuCEMe assay

For monocultures, cell lines were seeded in Thermo Scientific Nunc MicroWell white 96-well optical-bottom plates in 200 µL of their indicated culture media for 24 hours prior to analysis. Equal number of cells were seeded at 50 000 cells/well with four experimental conditions (vehicle, 2DG, OG and the combination) for each experimental group. 2DG and OG were added from a 10X stock and were used at a final concentration of 100 mM and 1 µM respectively. For luciferase-expressing cells, 150 µg/mL of D-Luciferin was added immediately after the addition of the metabolic inhibitors, and luminescence readout was performed using GloMax® Explorer Multimode Microplate Reader (Promega) for 30 minutes. For cells not expressing luciferase, ATP was measured using CellTiter Glo 2.0 according to the manufacturer’s instructions, after 30 minutes of exposure to the metabolic inhibitors.

For cocultures, MS5 (with or without adipogenic differentiation) and bone marrow macrophage cell line were cultured in Thermo Scientific Nunc MicroWell white 96-well optical-bottom plates in 200 µL of their indicated culture media for 24 hours. Then a PBS wash was performed prior to the addition of luciferase expressing cells (MLL-AF9 AML cells and MDA-MB-231 cells) cultured in their indicated culture media. Test cells were seeded at 50 000 cells/well (for each cell type) with four experimental conditions (vehicle, 2DG, OG and the combination) for each experimental group (monoculture vs coculture). After 24 hours of exposure, 2DG, OG and the combination was added to both experimental conditions followed by the addition of D-Luciferin. Luminescence readout was performed immediately using GloMax® Explorer Multimode Microplate Reader for 30 minutes.

For ex vivo analyses, spleen (for T cells), femurs, tibia and pelvis (for MLL-AF9 AML cells) of either leukemic mice or LPS rechallenged mice were harvested immediately following euthanasia. The bones were crushed using a mortar and pestle in phosphate-buffered saline (PBS) containing 2% FBS (flow buffer) and filtered using a 40 μm cell strainer. The spleens were homogenized using a plunger taken from a syringe and a 40 μm cell strainer. Red blood cells were lysed using ACK lysis buffer, after which CD8⁺ T cells were isolated by negative selection using the Mouse CD8⁺ T Cell Enrichment Kit, according to the manufacturer’s instructions. GFP^+^ MLL-AF9 AML cells were sorted from the bone marrow suspension using Cytek Aurora™ CS System. The cells were then immediately plated in Thermo Scientific Nunc MicroWell white 96-well optical-bottom plates at 50 000 cells/well in 200 µL of flow buffer. 2DG, OG and the combination was added followed by the addition of D-Luciferin. Luminescence readout was performed immediately using GloMax® Explorer Multimode Microplate Reader for 30 minutes.

### Modified LuCEMe assays

#### Measuring mitochondrial ATP using MTS-hFluc

For mitochondrial luciferase expression, HEK293T cells were seeded in 6-well plates at a density of 250 000 cells/well in complete RPMI1640 medium. 24 hours after seeding, the culture medium was replaced with fresh complete RPMI, and cells were transiently transfected using FuGENE HD Transfection Reagent. Transfection complexes were prepared in Opti-MEM I Reduced Serum Medium by combining 20 μL FuGENE HD Transfection Reagent with 9 μg of MTS-hFluc vector (2.2 μL reagent per μg DNA) according to the manufacturer’s recommendations. Complexes were incubated for 15 minutes at room temperature before being added dropwise to the cells. Cells were maintained under standard culture conditions for an additional 24 hours, after which they were harvested using trypsin-EDTA, counted, and reseeded into Thermo Scientific Nunc MicroWell white 96-well optical-bottom plates at 50 000 cells/well in 200 µL RPMI1640 medium to assess their mitochondrial luciferase expression by adding D-Luciferin and measuring luminescence levels. Then LuCEMe was performed according to the standard LuCEMe protocol mentioned above.

#### Glucose-free conditions and pyruvate modifications

For glucose deprivation experiments, cells were seeded under their standard culture conditions and allowed to adhere overnight. The following day, cells were washed once with PBS to remove residual glucose-containing medium before replacement with glucose-free RPMI 1640 medium. Cells were maintained in glucose-free medium for 24 hours, after which LuCEMe was performed as described above.

For pyruvate supplementation experiments, immediately prior to metabolic perturbation, sodium pyruvate was added to the designated wells from a 10X stock solution to achieve a final concentration of 100 mM (to saturate pyruvate availability throughout the 30-minute assay window, ensuring that mitochondrial substrate supply remained non-limiting during glycolytic inhibition). Following pyruvate supplementation, 2DG, OG or the combination was added to the appropriate wells, and ATP measurements were performed according to the standard LuCEMe protocol.

### Calculation of the metabolic parameters

Metabolic parameters were calculated from ATP measurements obtained under basal conditions and following treatment with 2DG, OG or the combination of both inhibitors. ATP values measured after treatment with both 2DG, and OG combined represent the assay background and were used for normalization of all calculated parameters. The following equations were used to derive metabolic dependencies and capacities:

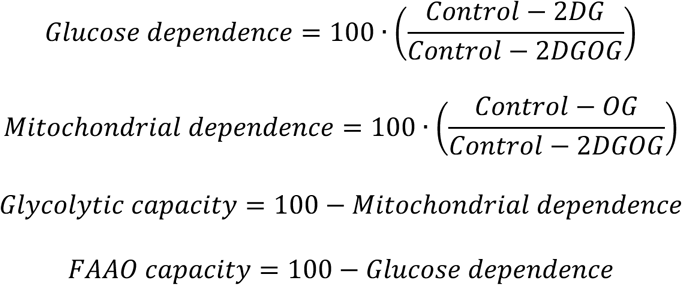

For experiments performed using pyruvate supplementation, additional metabolic parameters were calculated to distinguish glycolytic ATP production from glucose-derived mitochondrial ATP production. ATP measurements obtained in the presence of pyruvate following treatment with 2DG (2DG^pyruvate^), OG (OG^pyruvate^) or the combination of both inhibitors (2DGOG^pyruvate^) were incorporated into the following equations:

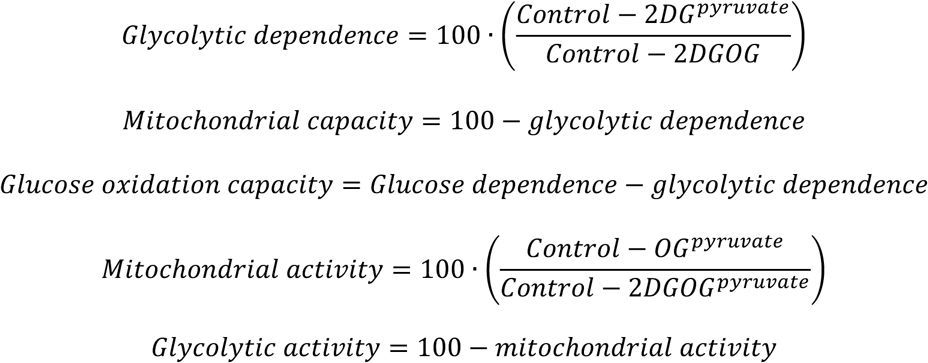

### Flow cytometry

Flow cytometry was performed to assess macrophage polarization, ^CD3+CD8+^T cell activation and exhaustion, and lipid peroxidation in AML cells. Unless otherwise indicated, cells were washed with flow buffer incubated with fluorophore-conjugated antibodies for 20-30 min at 4°C protected from light, washed, and resuspended in flow buffer for acquisition.

For macrophage polarization experiments, cells were stained with antibodies against CD80 and CD206 to identify pro-inflammatory and anti-inflammatory polarization states, respectively. Samples were acquired on a BD FACSVerse Flow Cytometer and analyzed using FlowJo software.

For ex vivo ^CD3+CD8+^T cell analyses, activation was assessed by staining for CD44, Ly6C, and CD25, whereas T cell exhaustion was evaluated using an antibody against PD1. Samples were acquired on a Cytek Aurora Spectral Flow Cytometer and analyzed using FlowJo.

Lipid peroxidation in MLL-AF9 AML cells isolated from mice at the time of maximal response to chemotherapy was quantified using the lipid peroxidation probe C11-BODIPY 581/591, according to the manufacturer’s instructions.

### SCENITH Analysis

For SCENITH analysis, MLL-AF9 AML cells were incubated with 2DG, OG, or the combination of both inhibitors at the same concentrations used for LuCEMe assay described above. Following 20 minutes of metabolic inhibition, puromycin was added to a final concentration of 10 μg/mL and cells were incubated for an additional 10 min to label newly synthesized proteins. Cells were subsequently harvested, fixed, and permeabilized prior to intracellular staining with an unconjugated anti-puromycin antibody, followed by the appropriate fluorophore-conjugated secondary antibody. Protein synthesis was quantified by flow cytometry using a BD FACSVerse Flow Cytometer, and data were analyzed using FlowJo. Relative protein synthesis under each inhibitor condition was used to calculate metabolic dependencies according to the original SCENITH methodology^5^.

### Extracellular Flux Analysis

For Seahorse assay, the OCR and ECAR were measured using a Seahorse XF96 analyzer (Seahorse Bioscience, Agilent) at 37 °C. MLL-AF9 AML cells were seeded at 50 000 cells/well in 180 μL XF RPMI medium. For the MitoStress assay, XF RPMI medium was supplemented with 10 mM D-glucose and 2 mM L-glutamine. For the GlycoStress assay, XF RPMI medium was supplemented with 2 mM L-glutamine. Plates were incubated for 60 minutes at 37°C in a CO_2_-free incubator to stabilize the pH. Analyses were performed under basal conditions and after the following sequential injections: MitoStress :1 μM OG (Port A), 1 μM FCCP (Port B) and 0.5 μM antimycin A (Port C). GlycoStress assay: 10 mM glucose (Port A), 2.5 μM OG (Port B) and 50 mM 2DG (Port C). The Seahorse Wave Desktop software was used to design assays and for data analysis, and measurements were normalized to the number of cells per well.

## QUANTIFICATION AND STATISTICAL ANALYSIS

Data are presented as mean ± standard deviation (SD). Statistical analyses were performed using GraphPad Prism 5. Statistical significance between two groups was determined using a two-tailed unpaired Student’s *t*-test. Comparisons among multiple groups were analyzed using one-way or two-way ANOVA followed by Tukey’s or Dunnett’s multiple-comparison test, as appropriate. A value of *P* < 0.05 was considered statistically significant.

## Declaration of generative AI and AI-assisted technologies in the manuscript preparation process

During the preparation of this work, the authors used ChatGPT (OpenAI) to improve language and readability and refine the phrasing. The authors reviewed and edited the output as needed and take full responsibility for the content of the published article.

## Supplemental Figures

**Supplemental Figure 1.**
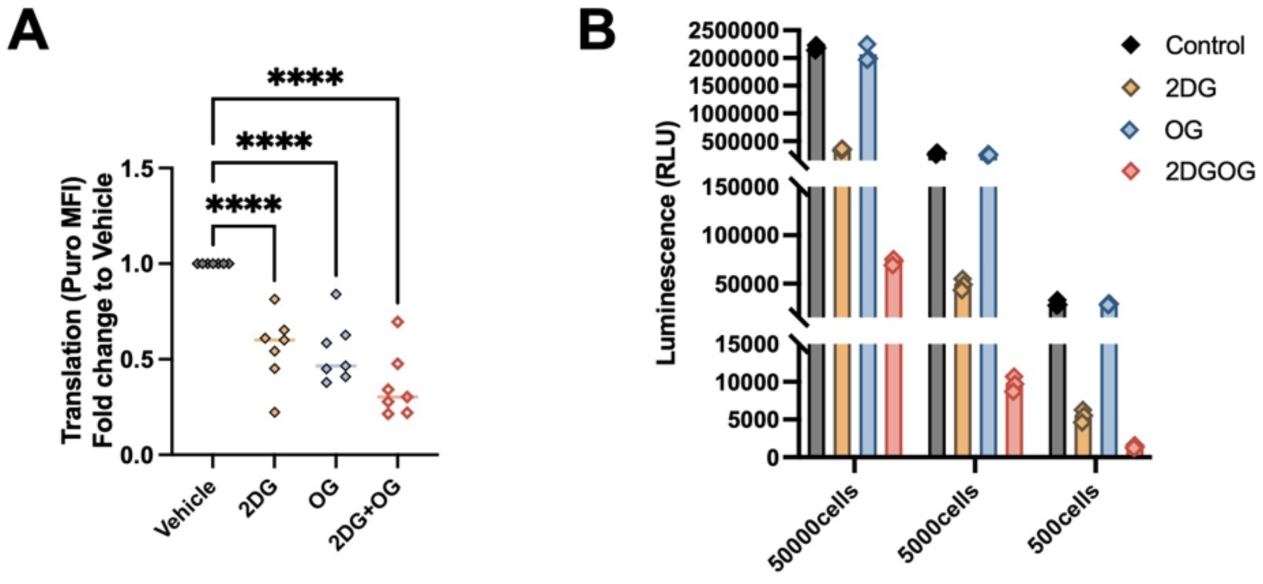
**(A)** Basal translation levels (anti-puromycin geometric mean) in MLL-AF9 AML cells after 30 minutes exposure to 2DG, OG and the combination. **(B)** ATP levels of AML cells seeded at different cell numbers measured using CellTiter Glo. Data are represented as mean ± SD. *p < 0.05, **p < 0.01, ***p < 0.001.

**Supplemental Figure 2.**
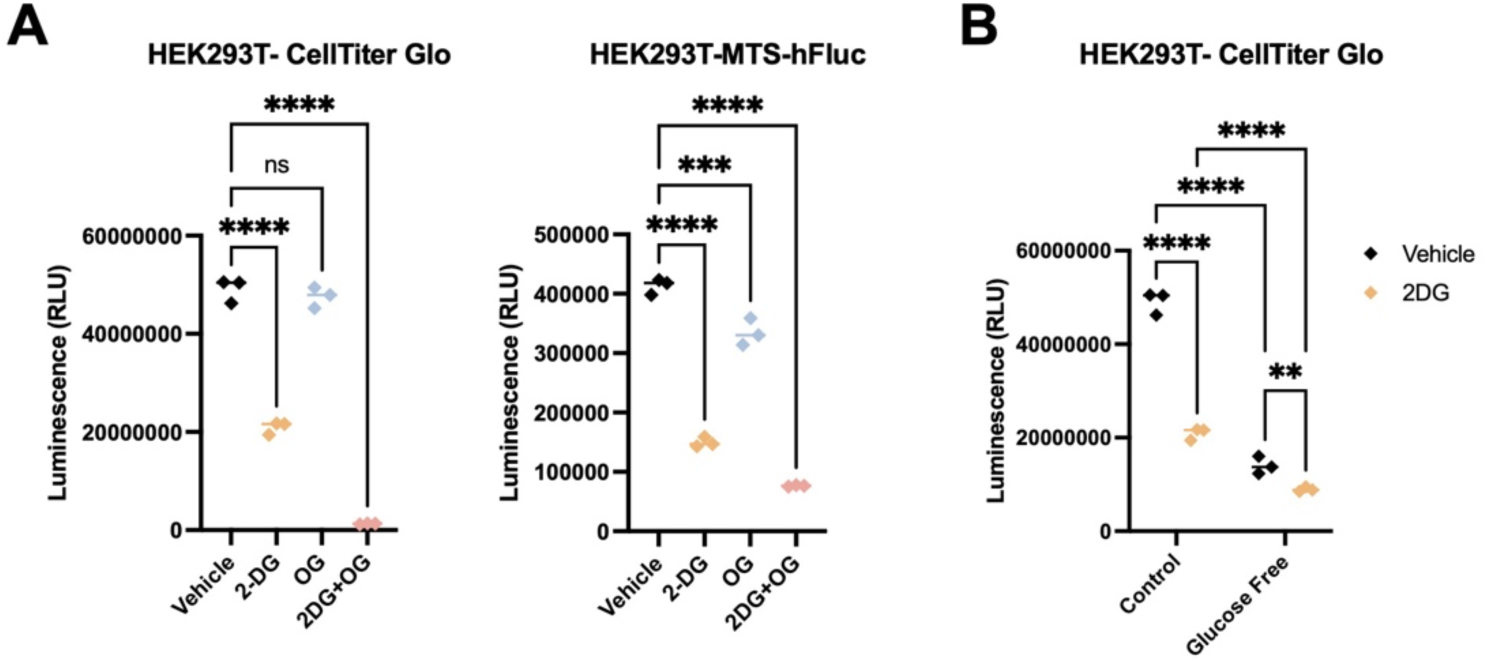
**(A)** Total ATP levels versus mitochondrial ATP levels measured using either CellTiter Glo or MTS-Fluc in HEK293T cells. **(B)** Total ATP levels in HEK293T cells after exposure to glucose-free conditions with or without 2DG treatment. Data are represented as mean ± SD.*p < 0.05, **p *< 0.01, ***p < 0.001*.

**Supplemental Figure 3.**
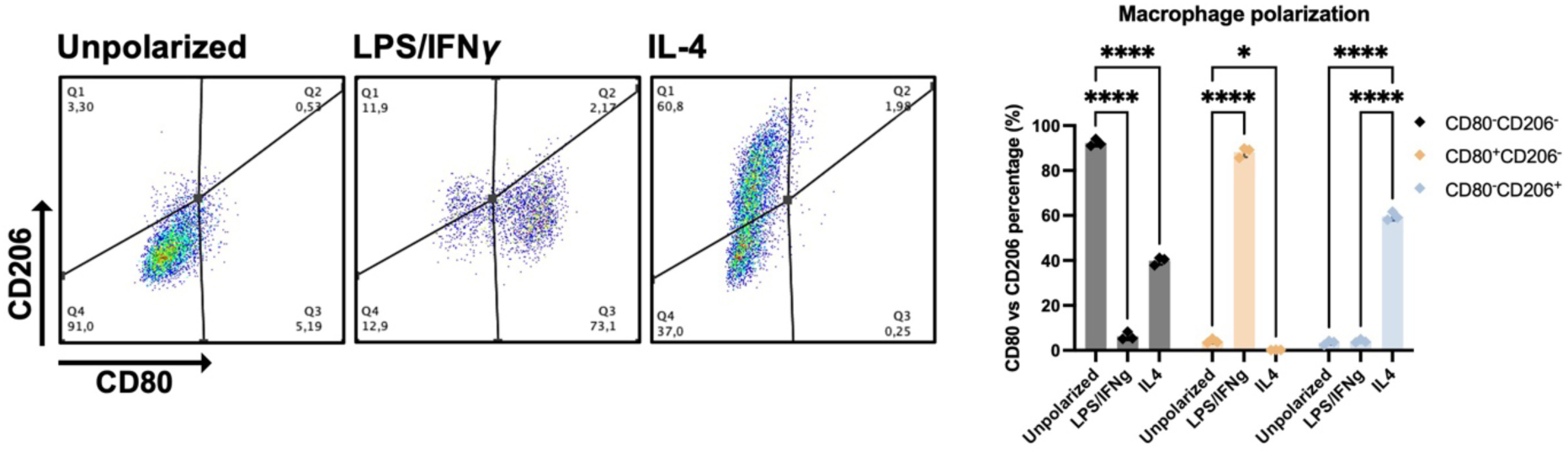
Flow cytometric analysis of CD80 versus CD206 levels in macrophages after 24 hours exposure to either LPS/IFNγ or IL-4. Data are represented as mean ± SD. *p < 0.05, **p < 0.01, ***p *< 0.001*.

**Supplemental Figure 4.**
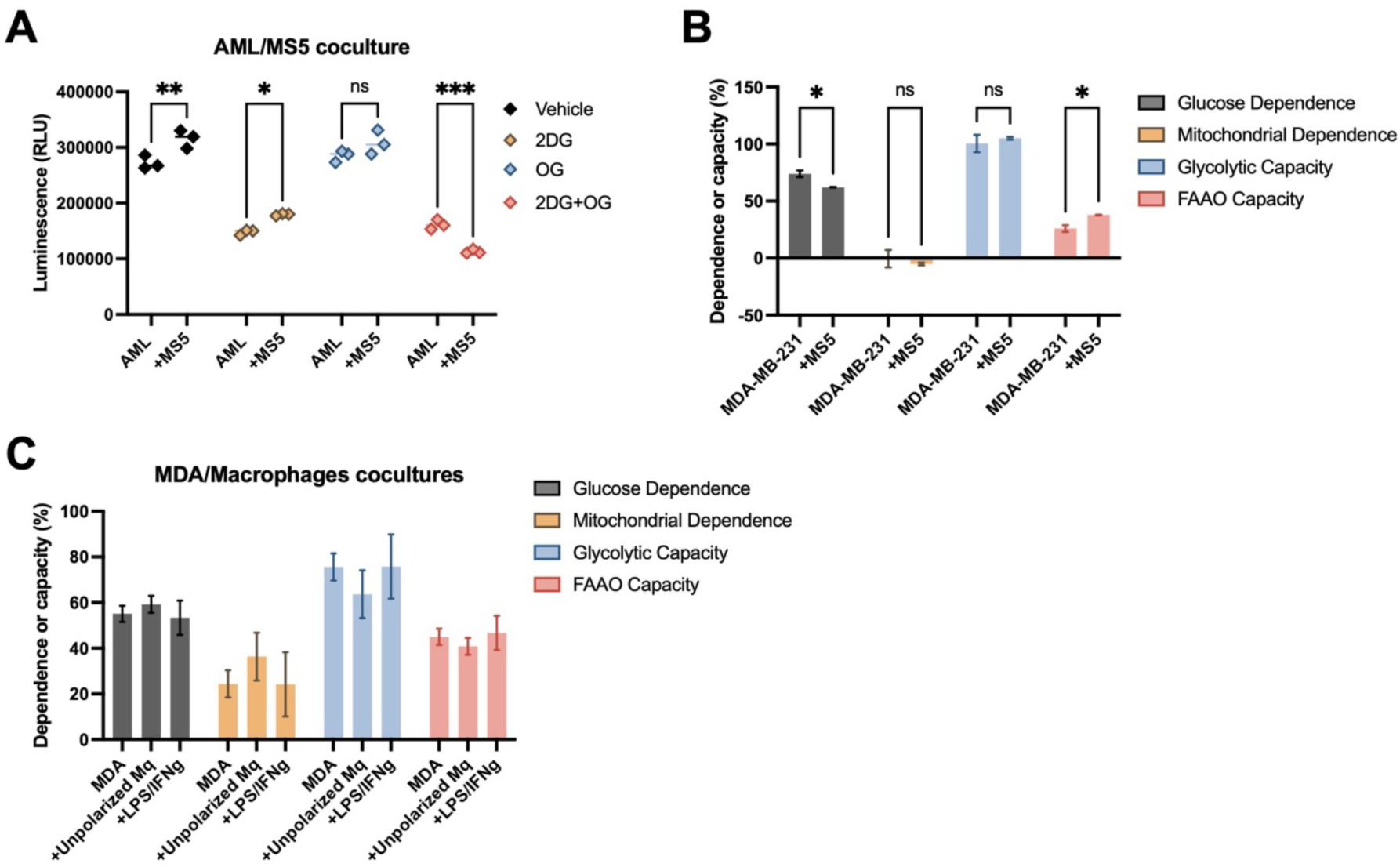
**(A)** Total ATP levels of MLL-AF9 AML cells in monoculture or coculture with MS5 cells after exposure to 2DG, OG and the combination. **(B)** Metabolic profile of MDA-MB-231 after coculture with MS5 cells for 24 hours, using LuCEMe. **(C)** Metabolic profile of MDA-MB-231 after coculture with macrophages with or without prior polarization with LPS/IFNγ, using LuCEMe. Data are represented as mean ± SD. *p < 0.05, **p < 0.01, ***p < 0.001.

## References

1. Pavlova, N.N., and Thompson, C.B. (2016). The Emerging Hallmarks of Cancer Metabolism. Cell Metab 23, 27–47. 10.1016/j.cmet.2015.12.006.

2. Rodriguez-Zabala, M., Ramakrishnan, R., Reinbach, K., Ghosh, S., Oburoglu, L., Falqués-Costa, A., Bellamkonda, K., Ehinger, M., Peña-Martínez, P., Puente-Moncada, N., et al. (2023). Combined GLUT1 and OXPHOS inhibition eliminates acute myeloid leukemia cells by restraining their metabolic plasticity. Blood Adv 7, 5382–5395. 10.1182/bloodadvances.2023009967.

3. Yoo, I., Ahn, I., Lee, J., and Lee, N. (2024). Extracellular flux assay (Seahorse assay): Diverse applications in metabolic research across biological disciplines. Mol Cells 47, 100095. 10.1016/j.mocell.2024.100095.

4. Desousa, B.R., Kim, K.K.O., Jones, A.E., Ball, A.B., Hsieh, W.Y., Swain, P., Morrow, D.H., Brownstein, A.J., Ferrick, D.A., Shirihai, O.S., et al. (2023). Calculation of ATP production rates using the Seahorse XF Analyzer. EMBO Rep 24, e56380. 10.15252/embr.202256380.

5. Argüello, R.J., Combes, A.J., Char, R., Gigan, J.P., Baaziz, A.I., Bousiquot, E., Camosseto, V., Samad, B., Tsui, J., Yan, P., et al. (2020). SCENITH: A Flow Cytometry-Based Method to Functionally Profile Energy Metabolism with Single-Cell Resolution. Cell Metab 32, 1063–1075. 10.1016/j.cmet.2020.11.007.

6. Xi, H., Kurtoglu, M., Liu, H., Wangpaichitr, M., You, M., Liu, X., Savaraj, N., and Lampidis, T.J. (2011). 2-Deoxy-d-glucose activates autophagy via endoplasmic reticulum stress rather than ATP depletion. Cancer Chemother Pharmacol 67, 899–910. 10.1007/s00280-010-1391-0.

7. Fessler, E., Eckl, E.-M., Schmitt, S., Mancilla, I.A., Meyer-Bender, M.F., Hanf, M., Philippou-Massier, J., Krebs, S., Zischka, H., and Jae, L.T. (2020). A pathway coordinated by DELE1 relays mitochondrial stress to the cytosol. Nature 579, 433–437. 10.1038/s41586-020-2076-4.

8. Huang, H.-L., Hsing, H.-W., Lai, T.-C., Chen, Y.-W., Lee, T.-R., Chan, H.-T., Lyu, P.-C., Wu, C.-L., Lu, Y.-C., Lin, S.-T., et al. (2010). Trypsin-induced proteome alteration during cell subculture in mammalian cells. Journal Biomed Sci 17, 36. 10.1186/1423-0127-17-36.

9. Danhier, P., Copetti, T., De Preter, G., Leveque, P., Feron, O., Jordan, B.F., Sonveaux, P., and Gallez, B. (2013). Influence of Cell Detachment on the Respiration Rate of Tumor and Endothelial Cells. PLoS One 8, e53324. 10.1371/journal.pone.0053324.

10. Ulrich, F. (1976). Effects of trypsin on protein synthesis in macrophages. Exp Cell Res 101, 267–277. 10.1016/0014-4827(76)90377-3.

11. van Gastel, N., Spinelli, J.B., Sharda, A., Schajnovitz, A., Baryawno, N., Rhee, C., Oki, T., Grace, E., Soled, H.J., Milosevic, J., et al. (2020). Induction of a Timed Metabolic Collapse to Overcome Cancer Chemoresistance. Cell Metab 32, 391–403. 10.1016/j.cmet.2020.07.009.

12. Farge, T., Saland, E., de Toni, F., Aroua, N., Hosseini, M., Perry, R., Bosc, C., Sugita, M., Stuani, L., Fraisse, M., et al. (2017). Chemotherapy-Resistant Human Acute Myeloid Leukemia Cells Are Not Enriched for Leukemic Stem Cells but Require Oxidative Metabolism. Cancer Discov 7, 716–735. 10.1158/2159-8290.CD-16-0441.

13. Maldonado, E.N., DeHart, D.N., Patnaik, J., Klatt, S.C., Gooz, M.B., and Lemasters, J.J. (2016). ATP/ADP Turnover and Import of Glycolytic ATP into Mitochondria in Cancer Cells Is Independent of the Adenine Nucleotide Translocator. J Biol Chem 291, 19642–19650. 10.1074/jbc.M116.734814.

14. Sawai, A., Taniguchi, T., Noguchi, K., Seike, T., Okahashi, N., Takaine, M., and Matsuda, F. (2025). ATP Supply from Cytosol to Mitochondria Is an Additional Role of Aerobic Glycolysis to Prevent Programmed Cell Death by Maintenance of Mitochondrial Membrane Potential. Metabolites 15, 461. 10.3390/metabo15070461.

15. Erdem, A., Kaye, S., Caligiore, F., Johanns, M., Leguay, F., Schuringa, J.J., Ito, K., Bommer, G., and van Gastel, N. (2025). Lactate dehydrogenase A-coupled NAD(+) regeneration is critical for acute myeloid leukemia cell survival. Cancer Metab 13, 22. 10.1186/s40170-025-00392-4.

16. Kuznetsov, A.V., Javadov, S., Sickinger, S., Frotschnig, S., and Grimm, M. (2015). H9c2 and HL-1 cells demonstrate distinct features of energy metabolism, mitochondrial function and sensitivity to hypoxia-reoxygenation. Biochim Biophys Acta 1853, 276–284. 10.1016/j.bbamcr.2014.11.015.

17. Raggi, F., Pelassa, S., Pierobon, D., Penco, F., Gattorno, M., Novelli, F., Eva, A., Varesio, L., Giovarelli, M., and Bosco, M.C. (2017). Regulation of Human Macrophage M1-M2 Polarization Balance by Hypoxia and the Triggering Receptor Expressed on Myeloid Cells-1. Front Immunol 8, 1097. 10.3389/fimmu.2017.01097.

18. Vrieling, F., van der Zande, H.J.P., Naus, B., Smeehuijzen, L., van Heck, J.I.P., Ignacio, B.J., Bonger, K.M., Van den Bossche, J., Kersten, S., and Stienstra, R. (2024). CENCAT enables immunometabolic profiling by measuring protein synthesis via bioorthogonal noncanonical amino acid tagging. Cell Rep Methods 4, 100883. 10.1016/j.crmeth.2024.100883.

19. Fuchs, A.L., Schiller, S.M., Keegan, W.J., Ammons, M.C.B., Eilers, B., Tripet, B., and Copié, V. (2019). Quantitative (1)H NMR Metabolomics Reveal Distinct Metabolic Adaptations in Human Macrophages Following Differential Activation. Metabolites 9. 10.3390/metabo9110248.

20. van Gastel, N., and Carmeliet, G. (2021). Metabolic regulation of skeletal cell fate and function in physiology and disease. Nat Metab 3, 11–20. 10.1038/s42255-020-00321-3.

21. Tirado, H.A., Balasundaram, N., Laaouimir, L., Erdem, A., and van Gastel, N. (2023). Metabolic crosstalk between stromal and malignant cells in the bone marrow niche. Bone Rep 18, 101669. 10.1016/j.bonr.2023.101669.

22. You, R., Hou, D., Wang, B., Liu, J., Wang, X., Xiao, Q., Pan, Z., Li, D., Feng, X., Kang, L., et al. (2021). Bone marrow microenvironment drives AML cell OXPHOS addiction and AMPK inhibition to resist chemotherapy. J Leukoc Biol 112, 299–311. 10.1002/jlb.6a0821-409rr.

23. Liu, W., Menoret, A., and Vella, A.T. (2017). Responses to LPS boost effector CD8 T-cell accumulation outside of signals 1 and 2. Cell Mol Immunol 14, 254–264. 10.1038/cmi.2015.69.

24. Cao, J., Liao, S., Zeng, F., Liao, Q., Luo, G., and Zhou, Y. (2023). Effects of altered glycolysis levels on CD8(+) T cell activation and function. Cell Death Dis 14, 407. 10.1038/s41419-023-05937-3.

25. Bengsch, B., Johnson, A.L., Kurachi, M., Odorizzi, P.M., Pauken, K.E., Attanasio, J., Stelekati, E., McLane, L.M., Paley, M.A., Delgoffe, G.M., and Wherry, E.J. (2016). Bioenergetic Insufficiencies Due to Metabolic Alterations Regulated by the Inhibitory Receptor PD-1 Are an Early Driver of CD8(+) T Cell Exhaustion. Immunity 45, 358–373. 10.1016/j.immuni.2016.07.008.

26. Wang, B., Tang, J., Zhou, Y., Huang, Q., Jiang, X., Zhang, F., Fan, S., Zhang, Q., Zhou, J., Wang, Y., et al. (2025). Abnormal metabolic activation of CD8(+) T cells correlates with poor prognosis in acute myeloid leukemia. J Transl Med 23, 791. 10.1186/s12967-025-06833-4.

27. Forte, D., García-Fernández, M., Sánchez-Aguilera, A., Stavropoulou, V., Fielding, C., Martín-Pérez, D., López, J.A., Costa, A.S.H., Tronci, L., Nikitopoulou, E., et al. (2020). Bone Marrow Mesenchymal Stem Cells Support Acute Myeloid Leukemia Bioenergetics and Enhance Antioxidant Defense and Escape from Chemotherapy. Cell Metab 32, 829–843. 10.1016/j.cmet.2020.09.001.

28. Yusuf, R.Z., Saez, B., Sharda, A., van Gastel, N., Yu, V.W.C., Baryawno, N., Scadden, E.W., Acharya, S., Chattophadhyay, S., Huang, C., et al. (2020). Aldehyde dehydrogenase 3a2 protects AML cells from oxidative death and the synthetic lethality of ferroptosis inducers. Blood 136, 1303–1316. 10.1182/blood.2019001808.

29. Sykes, D.B., Kfoury, Y.S., Mercier, F.E., Wawer, M.J., Law, J.M., Haynes, M.K., Lewis, T.A., Schajnovitz, A., Jain, E., Lee, D., et al. (2016). Inhibition of Dihydroorotate Dehydrogenase Overcomes Differentiation Blockade in Acute Myeloid Leukemia. Cell 167, 171–186. 10.1016/j.cell.2016.08.057.

30. van Gastel, N., Torrekens, S., Roberts, S.J., Moermans, K., Schrooten, J., Carmeliet, P., Luttun, A., Luyten, F.P., and Carmeliet, G. (2012). Engineering Vascularized Bone: Osteogenic and Proangiogenic Potential of Murine Periosteal Cells. Stem Cells 30, 2460–2471. 10.1002/stem.1210.

